# The limits of Bayesian estimates of divergence times in measurably evolving populations

**DOI:** 10.64898/2026.02.28.708707

**Authors:** Simona Ivanov, Samuel Fosse, Mario dos Reis, Sebastian Duchene

## Abstract

Bayesian inference of divergence times for extant species using molecular data is an unconventional statistical problem: Divergence times and molecular rates are confounded, and only their product, the molecular branch length, is statistically identifiable. This means we must use priors on times and rates to break the identifiability problem. As a consequence, there is a lower bound in the uncertainty that can be attained under infinite data for estimates of evolutionary timescales using the molecular clock. With infinite data (i.e., an infinite number of sites and loci in the alignment) uncertainty in ages of nodes in phylogenies increases proportionally with their mean age, such that older nodes have higher uncertainty than younger nodes. On the other hand, if extinct taxa are present in the phylogeny, and if their sampling times are known (i.e., ‘heterochronous’ data), then times and rates are identifiable and uncertainties of inferred times and rates go to zero with infinite data. However, in real heterochronous datasets (such as viruses and bacteria), alignments tend to be small and how much uncertainty is present and how it can be reduced as a function of data size are questions that have not been explored. This is clearly important for our understanding of the tempo and mode of microbial evolution using the molecular clock. Here we conducted extensive simulation experiments and analyses of empirical data to develop the infinite-sites theory for heterochronous data. Contrary to expectations, we find that uncertainty in ages of internal nodes scales positively with the distance to their closest tip with known age (i.e., calibration age), not their absolute age. Our results also demonstrate that estimation uncertainty decreases with calibration age more slowly in datasets with more, rather than fewer site patterns, although overall uncertainty is lower in the former. Our statistical framework establishes the minimum uncertainty that can be attained with perfect calibrations and sequence data that are effectively infinitely informative. Finally, we discuss the implications for viral sequence datasets. In a vast majority of cases viral data from outbreaks is not sufficiently informative to display infinite-sites behaviour and thus all estimates of evolutionary timescales will be associated with a degree of uncertainty that will depend on the size of the dataset, its information content, and the complexity of the model. We anticipate that our framework is useful to determine such theoretical limits in empirical analyses of microbial outbreaks.

**Significance statement:** Genetic sequences are routinely used to date the origins of outbreaks and the divergence of viral and bacterial lineages, but every such date carries uncertainty, and it has been unclear how much of that uncertainty more sequence data can remove. We show that for pathogens sampled repeatedly over time, precision depends not on how old an event is but on how close it lies to the nearest sample of known date, and that even large influenza and hepatitis B virus datasets fall far short of the precision that is theoretically attainable. This provides a simple diagnostic for judging how much confidence a molecular date deserves, and whether sequencing more genomes would improve its precision.

## 1 Introduction

The molecular clock states that the evolutionary rate of a protein or DNA molecule is roughly constant over time (Zuckerkandl, 1987). It has proven essential for understanding the timescale over which organisms evolve. In the case of infectious microbes, estimates of their time of emergence are largely based on evidence from the molecular clock (Boni et al., 2020, Durrant et al., 2024, Holmes et al., 2016).

There exists a range of molecular clock models that describe the evolutionary rate as a statistical process (Bromham et al., 2018, Ho and Duchêne, 2014). In this context, the ‘evolutionary rate’ refers to the combination of the instantaneous mutation rate and the long-term substitution rate (the rate at which mutations are fixed). Importantly, molecular sequence data on their own only convey information about molecular distance, such that all molecular clock models require additional calibration information about time or about the evolutionary rate. For example, the known time of divergence between two lineages or a previous estimate of the evolutionary rate of a closely related organism can be used as molecular clock calibrations (reviewed by Mello and Schrago, 2024).

Many microbes, notably viruses and bacteria, evolve sufficiently quickly that sequence data collected over the course of an outbreak (months or years) capture a measurable amount of evolutionary change. Such data, sometimes referred to as heterochronous data, are said to have been sampled from a ‘measurably evolving population’ (Biek et al., 2015, Drummond et al., 2003), and the sequence sampling times themselves can be used for calibration (reviewed by Rieux and Balloux, 2016). Datasets that involve ancient DNA may also be treated as being sampled from a measurably evolving population, and thus the age of ancient samples may also provide a source of calibration information (Perri et al., 2021).

A fundamental question of molecular clock models is how the uncertainty in the resulting estimates is determined by information from the sequence data. In conventional statistical problems, uncertainty in parameter estimates goes to zero as information in the data goes to infinity. In molecular clock analysis, information content in datasets is a function of the number of sites and number of loci (or data partitions) in the alignment (Rannala and Yang, 2007, Yang and Rannala, 2006), and thus longer sequence datasets are expected to convey more information than those that are short (e.g. whole genomes vs a single gene). Indeed, whole genome data have been shown to improve estimates of evolutionary rates and timescales in pathogen phylogenetic analyses, compared to datasets that involve a short portion of the genome (Biek et al., 2015).

The infinite-sites theory of the molecular clock (not to be confused with the infinite-sites model of population genetics, which assumes that every new mutation in a genome occurs at a unique site and never recurs at the same location, see Gillespie, 2004) was developed by Yang and Rannala, 2006 and Rannala and Yang, 2007 to describe the uncertainty in Bayesian estimates of time and rate as a function of the number of sites and loci in the alignment. They showed that, for phylogenies of extant taxa with infinitely long alignments (i.e., with an infinite number of sites and loci in the alignment) the uncertainty of node age estimates is proportional to their mean age, meaning that estimates for the ages of nodes close to the root have higher uncertainty than younger nodes. The infinite-sites theory of the molecular clock can be assessed by inspecting a regression of the uncertainty in node ages, such as the 95% highest posterior density (HPD) interval as a function of the mean node age. The extent to which the points deviate from the regression line determines whether the data are infinitely informative (Rannala and Yang, 2007, Yang and Rannala, 2006), and the slope determines the rate at which uncertainty increases towards the root of the tree. The main practical consequence of the infinite-sites theory is that uncertainty in divergence time estimates has a lower limit (Dos Reis and Yang, 2013).

The infinite-sites theory has been immensely useful for interpreting evolutionary timescales of many organisms (e.g. Álvarez-Carretero et al. 2022, Dos Reis et al. 2015). On the other hand, its behaviour for measurably evolving populations has not been explored. It is a particularly important point given that molecular evidence is essential for outbreak investigations. Our understanding of the time of origin of the 2013-2016 West African Ebola outbreak (Holmes et al., 2016), the COVID-19 pandemic (Boni et al., 2020), the ongoing global MPOX outbreaks (Parker et al., 2025), HIV-1 in humans (Korber et al., 2000), and of pandemic influenza viruses (Smith et al., 2009), are all grounded on the molecular clock.

Viral and bacterial genomes evolve over shorter timescales and at much higher rates than their eukaryotic counterparts. The Ebola virus, from the 2013-2016 outbreak, had an average evolutionary rate of 1.2 *×* 10*^−^*^3^ nucleotide substitutions/site/year (subs/site/year) (Holmes et al., 2016). When measured throughout a similar time frame, MERS-CoV had a mean rate of 7.8 *×* 10*^−^*^4^ subs/site/year (Boni et al., 2020). Bacteria tend to evolve more slowly than viruses, from 10*^−^*^6^ to 10*^−^*^8^ subs/site/year (Duchêne et al., 2016b). For comparison, evolutionary rates of eukaryotic organisms are orders of magnitude lower, ranging from 10*^−^*^8^ to 10*^−^*^10^ subs/site/year (see Fig. 1 in Holmes 2010).

**Figure 1:**
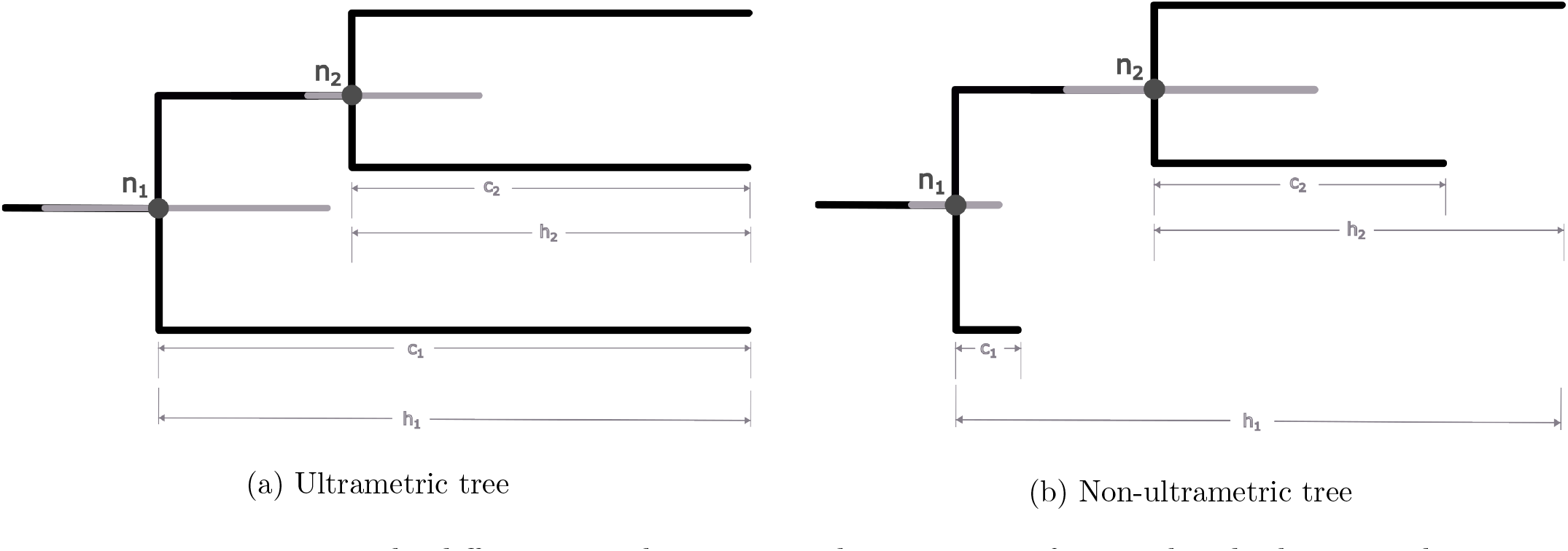
Here we can see the differences in the anticipated uncertainty of internal nodes between ultrametric and non-ultrametric trees. We use the *h*_1_ and *h*_2_ metrics to represent the heights of nodes 1 and 2 (*n*_1_ and *n*_2_) respectively, and *c*_1_ and *c*_2_ denote the distance to the closest tip (leaf node). In the ultrametric tree in Fig. 1a, it is evident that as *h*_1_ *> h*_2_, the uncertainty in the estimate (e.g. the 95% HPD), represented as the grey bar, of *n*_1_ is wider than that of *n*_2_. This is in line with the predictions of infinite-sites theory from previous studies, which predict a linear increase in divergence time uncertainty with node age for ultrametric trees, under the condition that the dataset is highly informative and that there is no additional calibration information. In contrast, we can observe that in Fig. 1b, for a non-ultrametric tree, this relationship does not necessarily hold. Although *h*_1_ *> h*_2_, the credible interval of *n*_2_ is actually wider than that of *n*_1_. If we focus on the distance to the closest tip of *n*_1_ and *n*_2_, represented as *c*_1_ and *c*_2_ respectively, we see that nodes that are closer to genome samples (tips of the tree) have lower uncertainties than those that are further away. We hypothesise that this occurs because the tips act as a lower bound that is not the same for all internal nodes in non-ultrametric trees.

In principle, the serially sampled nature of many microbial datasets means that their evolutionary rates and times are identifiable, and that the variance in their estimates would go to zero with infinitely informative data. However, real microbial datasets are relatively small in terms of the number of sites, and usually alignments are divided into one or just a few partitions, so that the data are much too small to observe such convergence to zero variance. This raises the question of how uncertainty in divergence time estimates scales with increasing numbers of sites in these datasets.

In this study, we investigate the rate at which uncertainty in time estimates in measurably evolving populations decreases as a function of data size. We focus on the Bayesian framework because uncertainty is well-characterised and a natural outcome of the inference, and because the original infinite-sites theory was developed under this framework (Yang and Rannala, 2006).

## 2 Results

### 2.1 Time to spill the information content

To study the behaviour of the infinite-sites theory in temporally structured data, we conducted a range of simulation experiments based on genome data from the H1N1 2009 influenza virus outbreak in North America. We analysed empirical data collected by Hedge et al., 2013 between January and June, August, or December (i.e. three cumulative datasets spanning January to June, January to August, and January to December, which we refer to as June, August, and December, respectively) using a Bayesian phylogenetic framework implemented in BEAST2 v2.7.7 (Bouckaert et al., 2019). We fixed the tree topology to the maximum likelihood estimate, and we set a HKY+Γ_4_ substitution model, a strict molecular clock and an exponential growth coalescent tree prior. We then summarised the ages of the nodes of the trees by calculating their 95% HPD intervals.

Strikingly, our initial analyses revealed no relationship between the absolute node age and their HPD interval width. We attribute this to the different nature of calibration information in ultrametric trees (i.e. those for which all samples were drawn at the same time) compared to those that are non-ultrametric (shown on Fig. 1a). In ultrametric trees the ages of all internal nodes must fall within the age of the root node and that of the tips. In contrast, for non-ultrametric trees, where samples have been collected over time as is the case for measurably evolving populations (Drummond et al., 2003), the ages of internal nodes fall between the age of the root node and that of their closest tip. For such time-structured data we considered the relationship between the node age relative to the age of the closest descendant tip (effectively the distance to the calibration, referred to here as the ‘tip-calibration’) and their associated uncertainty. In Fig. 1b we illustrate our argument: in (a) genome samples are all taken at the same time, and we observe a clear increase in HPD interval width as node age increases. In contrast, in (b) the HPD interval widths depend on the distance of the internal node to the sampled genome (tip or ‘leaf’ node).

In Fig. 2 we show the infinite-sites plot for a simulated dataset using the absolute node age or the node age relative to the closest tip-calibration. This demonstrates that in non-ultrametric trees, some internal nodes that appear towards the root of the tree, may appear to have a lower divergence time uncertainty than more recently-sampled nodes. In contrast, using the distance to the closest tip-calibration results in a nearly linear relationship between the two variables.

**Figure 2:**
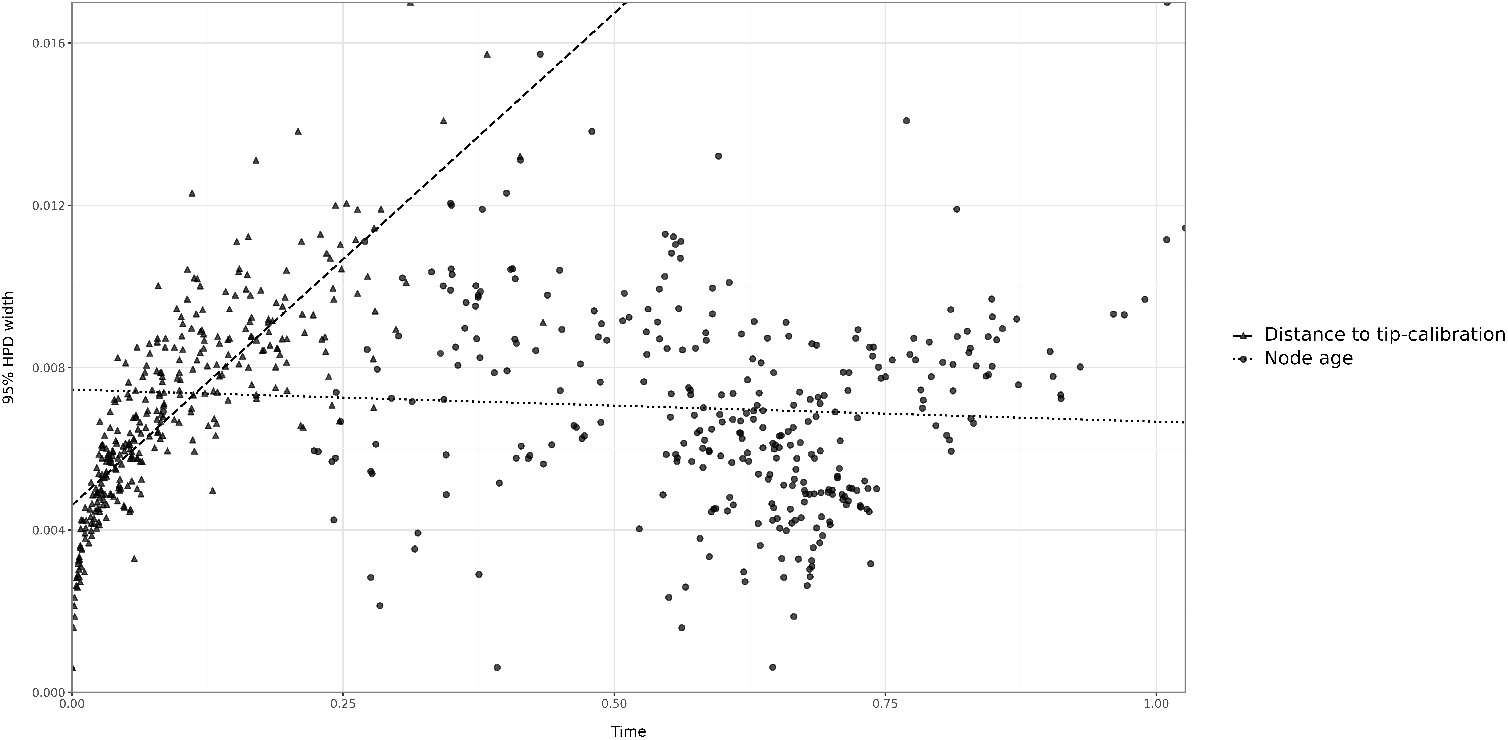
A comparison of the relationship between 95% HPD width and node age vs. distance to the closest tip-calibration. It is evident from the distribution of the points that the triangular points (i.e. node distance to tip-calibration vs. 95% HPD) shows a positive correlation, whereas the solid circles (i.e. node age vs 95% HPD) have no obvious relationship.

The goal of our simulations was to generate datasets with different degrees of information content. That is, those for which the sequence data provided little to no information about divergence times, and those that could be considered ‘infinitely informative’ (i.e. when we have all the information we could possibly have about the mutations occurring in the genome). The information content of a sequence alignment is typically determined via the sequence length (Yang and Rannala, 2012), and the number of unique site patterns (Lewis et al., 2016). However, it is important to note that information content is also relative to the size of the dataset (the number of taxa). In the context of phylogenetics, a given alignment length and number of unique site patterns will tend to provide higher resolution for a small, relative to a large number of taxa.

Given a fixed number of taxa and an evolutionary timescale, the number of unique site patterns is a function of the tree length (i.e. the total number of expected substitutions per site in a phylogenetic tree) and the sequence length, where longer sequences result in more opportunities for accruing mutations. In a sequence alignment, a unique site pattern refers to one arrangement of aligned bases that may or may not be repeated. In our simulation study we opted to vary the former, because it is more practical for computational purposes. Note, however, that in Bayesian and maximum likelihood inference, all sites in the alignment are informative, even if they are conserved.

To generate our simulated sequence alignments, we used the summary trees from analyses of the empirical data described above for each month (the full procedure is summarised schematically in Fig. S2 of the Supplemental Material). In order to achieve comparable numbers of unique site patterns across the three months (June, August, and December), we scaled the total length of each tree to 4 *×* 10*^−^*^4^, 5 *×* 10*^−^*^3^, and 2 subs/site (note that such scaling of the tree is similar to multiplying the branch lengths of the summary trees in units of time by an evolutionary rate). We simulated sequence evolution under alignments of 200,000 nucleotides and a HKY+Γ_4_ substitution model, which yielded around 80, 800, and 95,000 unique site patterns, for each tree length respectively. The simulation framework follows our analysis of the empirical data, where the influenza viral genomes taken from the middle of the outbreak, in August, contained on average around 800 unique site patterns (the mid-value of our simulations).

In Fig. 3, we show that the distributions of the number of unique site patterns are relatively similar across months for every tree length. However, it is important to note that as tree length increases, there is less overlap between different months, which can be clearly seen in Fig. 3c. We conjecture that this occurs because the datasets for later months have more sampled taxa, and thus their phylogenetic trees have more branches than those for earlier months. This, in turn, means that the average branch length for the datasets from these later months will be shorter and effectively allow us to observe more intermediate mutations, resulting in an overall greater number of unique site patterns (Duchêne et al., 2015, Hendy and Penny, 1989, Magallón, 2010, Stevenson et al., 2023).

**Figure 3:**
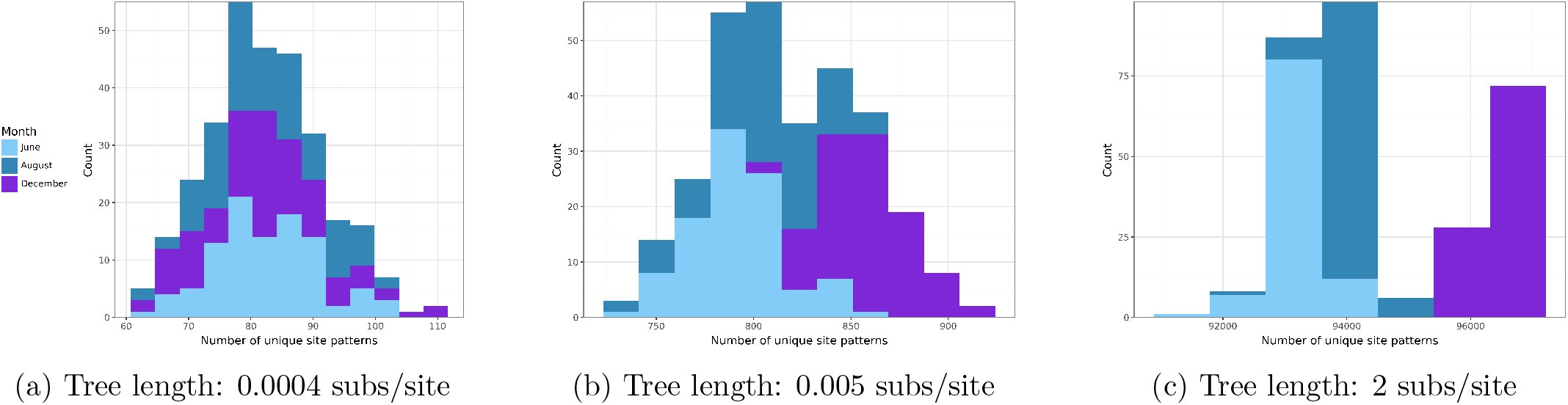
The distributions of the number of site patterns for different tree lengths across all three months after the empirical trees were scaled to the desired total tree lengths. Here the distributions are similar across different months. However, as the tree length increases, we observe that December still manages to accumulate more unique site patterns, despite our rescaling efforts. In the simulation for the month of December, we tend to have many short branches, as opposed to June, where we will see fewer branches that tend to be longer. Therefore, we hypothesise that this difference in the number of variable sites in later months could be due to the fact that we are able to observe many more intermediate mutations than in the earlier months.

### 2.2 To fix the topology or not to fix?

A major advantage of Bayesian phylogenetic analyses is that uncertainty is naturally accounted for in the posterior of parameter estimates. However, expressing uncertainty in divergence times when the tree topology is also estimated remains a challenging problem (Berling et al., 2025, Heled and Bouckaert, 2013). The most common approach is to select the tree topology that has the highest product of node support from the posterior, known as the maximum clade-credibility tree (known as the MCC tree) or the tree with the highest joint probability (known as the maximum ‘a posteriori’ or MAP tree). The uncertainty in the ages of internal nodes of the summary tree (MCC or MAP) are constructed by compiling the ages of trees from the posterior for which the node exists. Thus, nodes with very high topological support will have HPD intervals grounded on more samples from the posterior than those with low topological support (Heled and Bouckaert, 2013), and for this reason ages of internal nodes that are poorly supported may be misleadingly precise (Rannala, 2016).

For our initial analyses we attempted to investigate infinite-sites behaviour when the topology is estimated, but most simulations with a low number of unique site patterns had several nodes with no associated annotations. Essentially, many of the nodes in the MCC trees were present in very few trees from the posterior or entirely absent, meaning a very low posterior branch support (Fig. 4a). Because of this result, we decided to fix the tree topology to that used to generate each dataset (the ‘true’ tree), in order to be able to incorporate low numbers of unique site patterns, which were necessary for our analysis of the infinite-sites theory. This is a conservative assumption, but one that was necessary in order to isolate the sources of uncertainty we are interested in, and thus our results represent a lower bound on the uncertainty attainable when the topology is also estimated.

**Figure 4:**
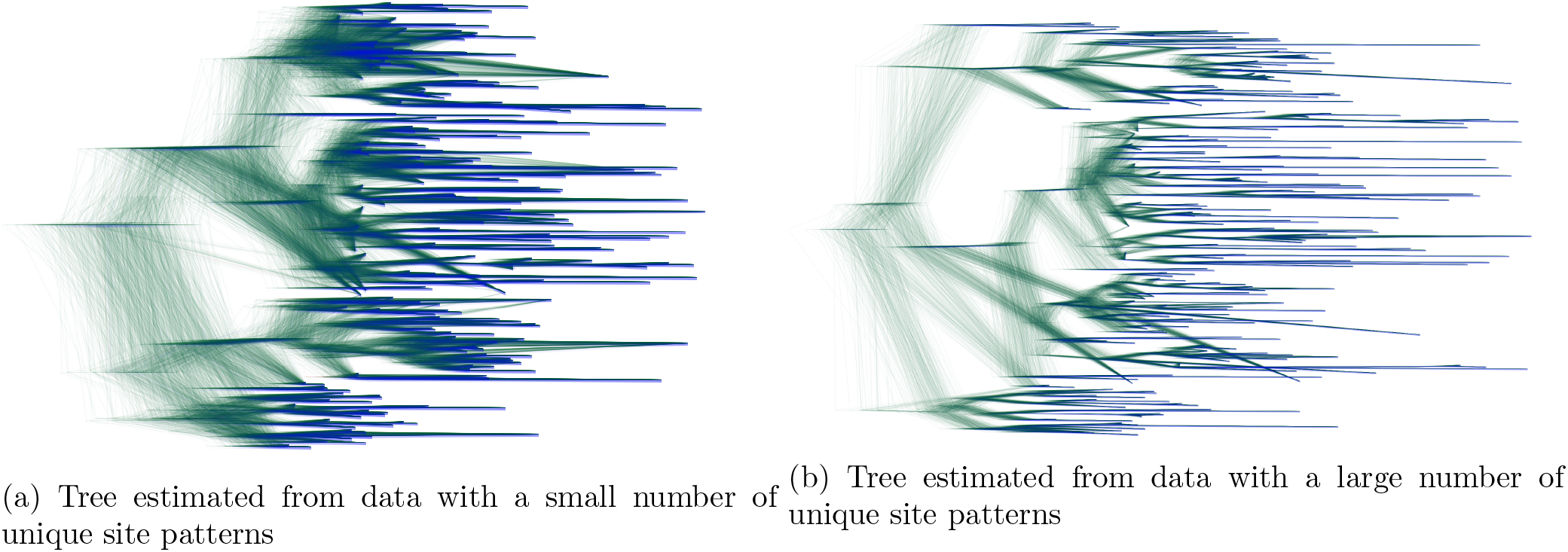
DensiTree figures for two datasets with the same number of taxa but estimated with (a) a small number of unique site patterns (788 unique site patterns) and (b) a large number of unique site patterns (2,198 unique site patterns). The cloud of lines represents individual trees sampled from the posterior in analyses where the tree topology was estimated. In (a) the cloud of trees is more fuzzy, meaning higher uncertainty in the topology and node heights, whereas in (b) the cloud of trees is more cohesive, consistent with lower uncertainty in the topology and node heights. The visualisations were made using DensiTree (Bouckaert, 2010).

### 2.3 Reaching for infinity

We fit a linear regression model for the HPD interval width of the age of each internal node as a function of their mean distance to their closest tip-calibration, for each simulated dataset. In Fig. 5 we show several such regressions, where we can clearly observe that the slope of the lines decreases, as tree length and the number of unique site patterns increase. It is also evident that the distribution of points become increasingly linear as we increase tree length (Fig. 6c). These results support the predictions of the infinite-sites theory: increasing information content results in a relationship between node-age uncertainty and distance to the nearest sampled genome that is increasingly linear, with a flatter slope, and with overall lower uncertainty.

**Figure 5:**
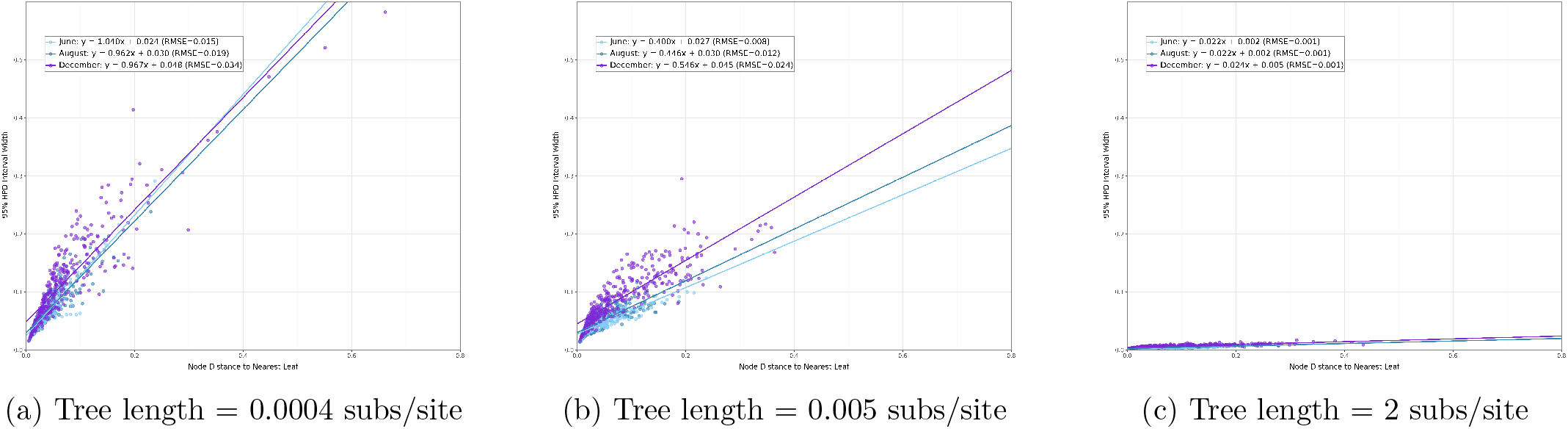
Infinite-sites plots for the three different tree lengths: 0.0004, 0.005, and 2 subs/site. As we increase the total tree length, the slope decreases, showing that as more site information is added, uncertainty increases at a slower pace and we can reach much lower overall amounts of uncertainty in our divergence time estimates. An interesting note is that data from the month of December has higher levels of uncertainty than the earlier months here, which we attribute to the fact that even though December maintains more unique site patterns the added information is not sufficient to balance the uncertainty imposed by the larger number of parameters that our model has to estimate (i.e. more taxa imply a larger parameter space).

**Figure 6:**
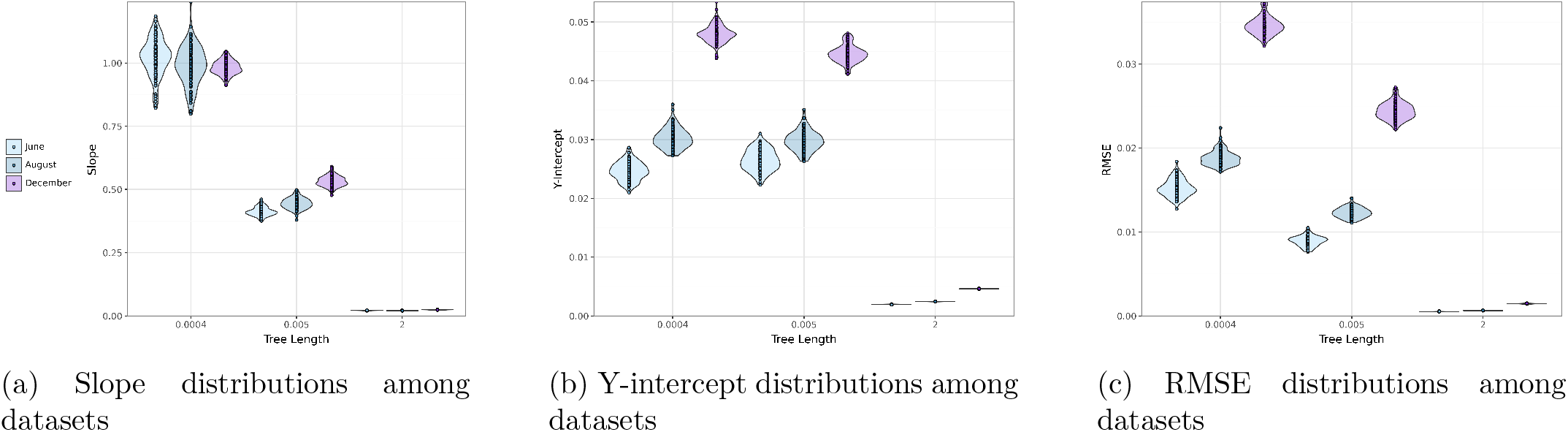
Distributions of linear regression statistics for simulation scenarios of months and tree lengths. Firstly, in Fig. 6a, we observe that as tree length increases, the slope decreases, demonstrating once again that the uncertainty in our estimates of divergence time does indeed decrease as we add more site information. We also observe that the y-intercept and the RMSE (root mean squared error) decrease as tree length increases (Figs. 6b and 6c), which also supports the predictions of infinite-sites theory, that with more site information, the observed relationship becomes more linear. Note that trees with increasing numbers of taxa (e.g. December compared to June) tend to have larger regression statistics, which likely occurs because they effectively contain more parameters (e.g. branch lengths) and thus require more sequence information.

In our simulations, attaining around 95,000 site patterns in the genome was needed for such infinite-sites behaviour, which illustrates that a theoretical limit of divergence time uncertainty can be reached if the data are asymptotically infinitely informative. In our simulated trees, we assumed that the age of each node would be informed by its closest child node and that its uncertainty scales with the distance between them. However, in simulated data, where we supposedly know the divergence times, it is possible that the closest leaf node of the parent is a shorter distance away from the node of interest than its own closest child. We investigated whether this phenomenon affects our simulations and found that it occurs in about 7-8% of our trees. We then updated the Python script used to find the closest leaf node of every internal one in order to find the minimum of the two distances described above and compared the infinite sites plots to the ones we had previously made. We did not observe any major differences in the relationship between node age and HPD width, but we have included the infinite sites plots that account for these cases in the Supplemental Material (Figs. S3–S5).

Importantly, the y-intercept of our regression was not strictly zero (Fig. 6b). This regression parameter represents the uncertainty in the age of an internal node whose distance to the closest tip is zero. The non-zero intercept occurs because this parameter corresponds to irreducible uncertainty from the estimate of the rate and the prior specified. This point is particularly relevant for genome samples drawn from an outbreak that emerged recently, where genome samples may have been collected immediately after divergence events, providing a low amount of information for the model to use to extrapolate divergence times. Indeed, estimates of the time of origin for outbreaks with intensive genome samples do not have zero uncertainty, even if samples were collected shortly after the origin of the outbreak, with SARS-CoV-2 and the 2013-2016 West African Ebola virus outbreaks being key examples (Boni et al., 2020, Holmes et al., 2016).

The infinite-sites plots between datasets from different months display some key patterns. Fig. 6 shows the resulting plots for the three months across tree lengths. We find that the slope, the RMSE, and the y-intercept all decrease with tree length, as expected. Importantly, when the tree length is shortest (0.0004 subs/site and lowest number of unique site patterns) or longest (2 subs/site and largest number of unique site patterns), we see very minor differences in the slope parameter for the three months (see also Fig. 6a). This likely occurs because in analyses of data with a small number of unique site patterns the prior is more influential with respect to the phylogenetic likelihood, compared to those with many site patterns. In the latter we expect that the phylogenetic likelihood will be overwhelmingly more influential than the prior and thus such datasets with many site patterns are expected to display infinite-sites behaviour. When there is an intermediate number of unique site patterns, as in our simulations with a tree length of 0.005 subs/site, we observe that datasets with more taxa (June has 164 taxa, while December has 328) have steeper slopes, and higher RMSE and y-intercepts. We attribute this to the fact that adding taxa in a phylogenetic analysis also means the addition of parameters, such as branch lengths and internal nodes, and thus a given number of unique site patterns conveys more information to a dataset with few taxa than to one with many taxa.

### 2.4 What about the clock model? Relax!

We have assumed so far a strict molecular clock, but many pathogens do not follow its assumption that all lineages evolve at the same rate, notably many viruses and bacteria display high among-lineage rate variation (Duchêne et al., 2016a,b). We explored the relaxed molecular clock model that assumes that branch rates are independent draws from a lognormal distribution (Drummond et al., 2006). This model has more parameters than the strict molecular clock model. For *s* taxa it has 1 +(2*s −* 2) = 2*s −* 1 more parameters than the strict clock: the variance of the lognormal rate, *σ*^2^, and 2*s −* 2 branch rates (Baele et al., 2014, Douglas et al., 2021, Rannala and Yang, 2007).

We anticipate the need for even more unique site patterns to reduce uncertainty in node ages, as we observed under the strict molecular clock model. Nevertheless, a key consideration is that infinite-sites theory was developed as a function of the number of sites and as a function of the number of loci (i.e, alignment partitions) (Zhu et al., 2015). The latter is important for relaxed molecular clocks: if we only have one partition, then the model is not fully identifiable and thus these analyses are not expected to yield complete infinite-sites behaviour.

As expected, under the same tree length and number of taxa, we observe higher uncertainty in the estimates with a relaxed clock model as opposed to those from the strict molecular clock (see Figs. 7b and 7c). However, it is noteworthy that when the number of unique site patterns is low, with a tree length of 0.0004 subs/site (Fig. 7a) the infinite-sites plots are nearly identical for both molecular clock models. This can be explained because information content in the sequence data is very low, such that the prior is more influential than the likelihood. In the relaxed molecular clock the default prior on the standard deviation of the branch rates in BEAST2, a parameter that determines the degree of among-lineage rate variation, is a Gamma distribution with shape=0.54 and rate=0.38 that places most weight on values close to zero, and thus penalises high rate variation among branches. As a result, this prior acts as a form of Bayesian regularisation (see Panchaksaram et al. 2025 and Ho et al. 2015 for the impact of the prior on rate variation in model selection). To investigate this pattern we conducted 20 replicates of each month-tree length combination, all of which confirmed the pattern shown in Fig. 7.

**Figure 7:**
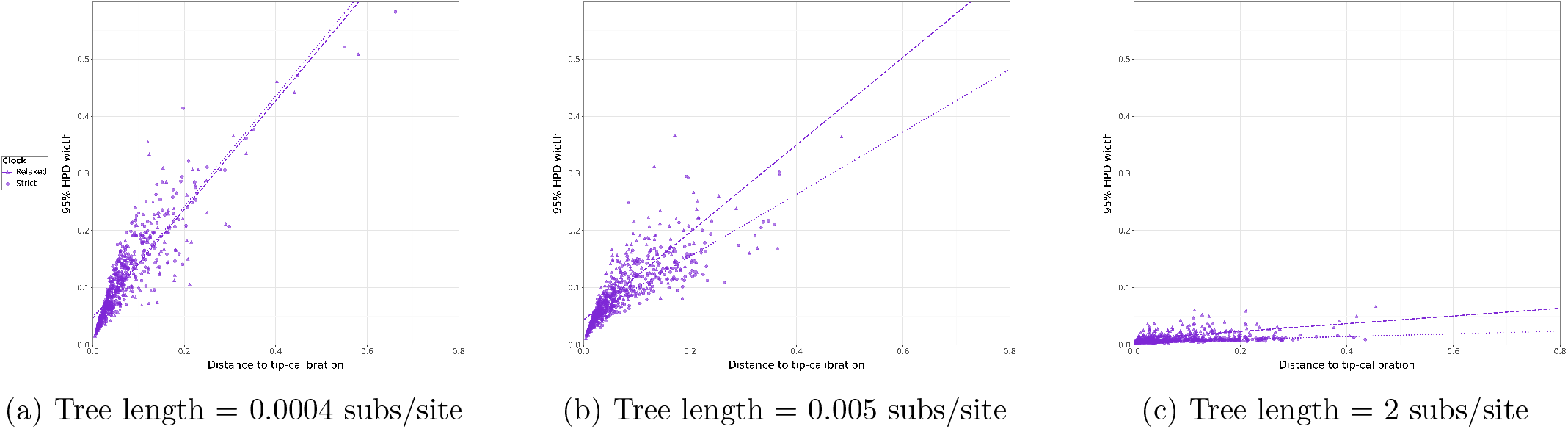
Here we compare infinite-sites plots, all derived from December datasets, but for different total tree lengths, to investigate how the choice of clock models affects the uncertainty in our estimates of divergence time. Overall, we see an increase in uncertainty for the relaxed clock case, as opposed to the strict clock, due to the fact that the relaxed clock model must estimate many more parameters and requires much more site information to do so.

### 2.5 It is a case study of flu and Hepatitis B (HBV)

We conducted analyses of empirical data from influenza virus from the 2009 pandemic sampled in North America from the start of the year until December (Hedge et al., 2013), as used in our simulations, and for a dataset of Hepatitis B virus (HBV) of 100 genomes that includes modern samples and those from over 10,000 years before present, as generated by Kocher et al. (2021) and curated by Weber et al. (2025). We chose these datasets because influenza virus has a reasonably constant sampling rate and is a widespread respiratory virus with a high evolutionary rate and an RNA genome (Duchêne et al., 2014). In contrast, HBV is a DNA virus that has been thought to co-diverge with humans for tens of thousands of years and tends to evolve more slowly than respiratory RNA viruses (Mühlemann et al., 2018), such that our case studies cover a range of expectations in virus evolution.

Interestingly, in this case we observe that influenza seems to be closer to infinite-sites behaviour than HBV, even though it has nearly half the number of unique site patterns (see Fig. 8). This result shows that infinite-sites behaviour is a combination of the evolutionary rate, the timescale and the number of calibrations. For example, note that HBV has been sampled over a period of more than 10,000 years, whereas the influenza virus used in this study contains samples from under one year. Moreover, the influenza dataset contains over three times as many sampled taxa as HBV. Thus, the higher levels of uncertainty seen in HBV as opposed to influenza, can be explained by the fact that there are fewer site patterns per taxon and the much longer time frame of the data.

**Figure 8:**
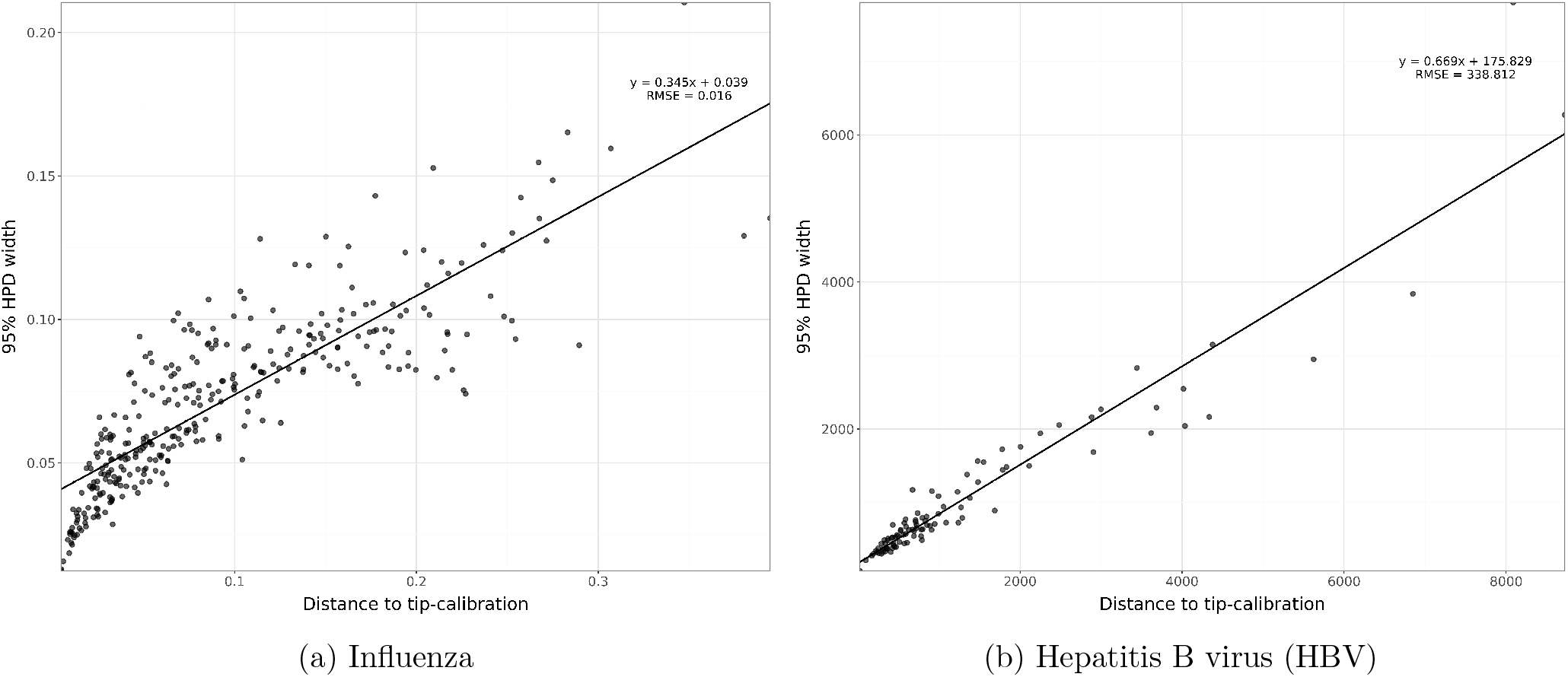
Here, we have infinite-sites plots for two empirical datasets: influenza virus from the 2009 H1N1 outbreak, sampled from early 2009 to December, which was also used as a basis for our simulation study, and 100 genomes of HBV. The sequence alignment used to generate the influenza plot contained about 1,500 unique site patterns, and that for the HBV contained approximately 2,100 unique site patterns. Note that the x- and y-axes are vastly different, which explains differences in regression coefficients and statistics.

## 3 Discussion

Our study describes the asymptotic behaviour of uncertainty in divergence times in the context of Bayesian phylogenetic analyses of time-structured data for which time-trees are non-ultrametric. Previous literature develops the infinite-sites theory, and in particular, its implications for datasets where time-trees are ultrametric (Dos Reis and Yang, 2013, Rannala and Yang, 2007). For heterochronous data that are typically collected from measurably evolving populations, we find that the infinite-sites plot (the regression of uncertainty in node ages as a function of their mean distance to the closest tip-calibration) is a valuable assessment of the information content in the data and the inherent uncertainty that can be expected. The slope, RMSE, and y-intercept provide important insights into our results.

The slope effectively quantifies the rate at which uncertainty in the age of internal nodes increases as their tip-calibrations become more distant. When sequence data are maximally informative, this parameter approaches zero, meaning that deep internal nodes that are distant from their tip-calibration have comparable precision to those that have a tip-calibration that is close in age. In this case the y-intercept will also be very low. For datasets that do not display such infinite-sites behaviour, the slope will tend to be larger, implying that long branch lengths are estimated with higher uncertainty than those that are short. Therefore, including tip-calibrations that are phylogenetically close to an internal node of interest and of a similar time frame are essential to improve the precision in the estimate of the node of interest. In our simulations of phylogenetic trees of length 4 *×* 10*^−^*^4^ subs/site, we obtained slopes of about 1.0, which implies a one-to-one association between uncertainty and mean distance to tip-calibration. For phylogenetic trees of length 2 subs/site, the association nearly disappears, with slope values that are over an order of magnitude lower. For our empirical data from influenza virus we found a slope of 0.345, about half of that of HBV at 0.669.

We have taken the tip-calibration of an internal node to be its closest descendant tip throughout. A tip that is not a descendant can also calibrate a node, by way of a path that runs from the node up to its parent and back down, and such a tip is sometimes nearer in time than any descendant. We investigated this case directly with a set of four-taxon simulations (Section 2 of the Supplemental Material). Both kinds of tip carry calibration information, and when the two are compared at matched distances the closest descendant carries the slightly larger effect. Where a non-descendant tip is much nearer in time than any descendant, however, it accounts for more of the uncertainty in the age of the node than the closest descendant does. We retain the closest descendant as the tip-calibration because it is the commonest pattern in the trees we analyse and the one most readily generated by coalescent and birth-death processes (Möller et al., 2018), and because identifying the tip that is closest in time requires the true node times, which are not available for empirical data. In practice we recommend comparing infinite-sites plots built on the closest descendant tip against plots built on the closest non-descendant tip, rather than relying on either alone.

In this study we assume that the sequence alignment is governed by a single genealogy, meaning that there is no recombination or reassortment. Analyses of some viruses and bacteria, and notably those that encompass deep evolutionary time frames may display substantial recombination or reassortment and therefore the effective alignment length is bounded by the non-recombinant segment size and not necessarily the whole genome length. Consequently, empirical datasets are likely far from displaying infinite-sites behaviour.

In our regression the y-intercept represents the theoretical minimum uncertainty that can be reached for an internal node whose mean estimated height is the same as its closest tip-calibration. Although it is not necessarily zero because tip-calibrations impose a minimum bound for the age of their parent node but not a maximum bound, we expect it to be very low for datasets with high information content. In our simulations, we found that this parameter could correspond to an uncertainty of about one to two weeks (1 year *×* 0.04 or 1 year *×* 0.02 for tree lengths of 4 *×* 10*^−^*^4^ subs/site to 2 subs/site; see Fig. 5). In our empirical datasets we see stark differences in this parameter, with 0.039 for influenza virus that implies a minimum uncertainty of about two weeks, while that for HBV was 175.8 years.

The RMSE measures how closely the uncertainty derived from the data follows a linear relationship as predicted by the original infinite-sites theory, with low values indicating that we can better predict uncertainty of node age estimates. These values will depend on the timescale of measurement, as the RMSE is the square root of the mean squared difference between prediction and truth, without normalisation. In our influenza virus data, with a timescale of about a year, we found a value of 0.016, whereas our HBV dataset, with a timescale of several thousands of years, had a value of 338.

Empirical analyses can benefit from the routine inspection of the infinite-sites plot. Modern outbreak investigations of viruses and bacteria generally consist of complete genome data, such that it is not possible to reduce uncertainty in estimates of node ages by sequencing longer genomes. In these cases, the infinite-sites plot can reveal the minimum uncertainty that we can expect in divergence events. Microbes with higher evolutionary rates will tend to produce more informative sequence datasets, for example the uncertainty in HBV, which has genome-wide evolutionary rates of the order of 10*^−^*^6^ subs/site/year, will attain an uncertainty of hundreds of years at best, whereas in influenza virus, with rates of around 10*^−^*^3^ subs/site/year, the uncertainty can be as low as a few weeks.

We note that the exact behaviour of the infinite-sites plot, however, depends on the information content in the data (sequence length and site patterns), the number of taxa, and their sampling time span. A sample taken from the middle of the H1N1 influenza epidemic, in the month of August, yields 807 unique site patterns (Hedge et al., 2013). Other rapidly-evolving RNA viruses, such as Ebola virus sampled in Sierra Leone during the 2013-2016 West African Ebola outbreak had only 28 site patterns (Gire et al., 2014, Stadler et al., 2014). Although Ebola virus has an evolutionary rate of about 2 *×* 10*^−^*^3^ subs/site/year (Gire et al., 2014), this localised introduction in Sierra Leone was sampled for less than a month, with insufficient evolutionary change to yield the same precision as an influenza virus outbreak unfolding over many months. Consequently, infinite-sites behaviour does not only depend on the organism (through its evolutionary rate and genome length), but also on the data themselves. Moreover, here we have only considered the situation where the data consist of a single partition, but the infinite-sites theory demonstrates that multiple partitions (i.e. loci) are necessary for the identifiability of relaxed molecular clocks (Zhu et al., 2015), a topic that warrants further work. We also considered a single parameterisation of the uncorrelated relaxed clock. Additive uncorrelated relaxed clocks offer a more realistic modelling approach (Didelot et al., 2021), in which the inferred number of substitutions along a branch is independent from the number of nodes along the path (Guindon, 2020). We anticipate that they will have a similar infinite-sites behaviour to the traditional uncorrelated relaxed clock as used here, although the exact degree of uncertainty may differ in magnitude.

An important conclusion we can draw from comparing empirical data and our simulations is that it is unlikely that a microbial molecular dataset sampled over a few months or years would exhibit true infinite-sites behaviour. Our simulations with the largest number of unique site patterns and which approach infinite-sites behaviour (slope approaching zero and very low y-intercept and RMSE) had 92,000 unique site patterns or more for a genome size of 200,000 nucleotides, for our simulations of the month of June. The corresponding evolutionary rate was about 0.12 subs/site/year, a value that is highly implausible given that the evolutionary rate scales negatively with genome size (Kuo et al., 2009, Lynch, 2010). For example, the MPOX virus has a similar genome size to our simulations, at around 197,000 nucleotides, but a much lower evolutionary rate, between 10*^−^*^6^ and 10*^−^*^4^ subs/site/year, depending on the genomic region (Parker et al., 2025).

Our findings support the view that the inclusion of ancient DNA samples is necessary for improving precision of divergence times, but their utility depends on their actual age and the microbe in question. A genome from variola virus from the 17*^th^* century narrowed down the origin of human clades in the decades preceding the pre-vaccination era (Duggan et al., 2016). However, samples from several thousand years ago were needed to detect any form of temporal signal for HBV (Duchêne et al., 2020, Kocher et al., 2021). The value of such ancient samples is that they are likely to increase the number of unique site patterns and they tend to act as tip-calibrations that are close to ancient nodes.

Our study has many interesting implications for ancient DNA analyses. We speculate that genomes with uncertainty in their sampling time, due to radio carbon dating, for example, also have a theoretical minimum uncertainty. Future studies could investigate how such uncertainty behaves in the context of the infinite-sites theory, particularly for ancient genome samples with associated priors on sampling times. We emphasise that the uncertainty might depend on the prior, the model and eventually calibration uncertainty that is used for tip-calibrations dated using radio carbon dating. It is also important to highlight that here we fixed the tree topology because the goal of our experiments is to assess precision of divergence time estimates in isolation. While integrating over uncertainty in the tree topology is important for phylodynamic analyses in practice (Fourment et al., 2025), it would be a confounding factor for disentangling uncertainty due to divergence time estimation, particularly for poorly supported nodes (Baele et al., 2025).

Finally, we note that here we focused on microbial data sampled through time, but our work should be informative on analyses of other measurably evolving populations such as ancient DNA studies (Bos et al., 2019), or studies of morphological datasets in which morphological characters have been scored for ancient fossil taxa (Ronquist et al., 2012). In all these cases the resulting time-trees are not ultrametric and the behaviour of time uncertainties should recapitulate some of the trends we describe here.

## 4 Materials and methods

In our simulation study we generated 900 summary trees under a strict molecular clock model, and 180 under a relaxed molecular clock model. These summary trees correspond to the fixed tree topology, with node heights scaled to their mean and with their uncertainty calculated from all trees in the posterior, similar to a maximum clade-credibility tree with node heights reported as the mean, but with a fixed topology. We scaled the empirical trees from three months of the 2009 H1N1 Influenza epidemic, studied by Hedge et al. (2013), to three different tree lengths, in order to simulate a range of unique site patterns. The three lengths used were 4 *×* 10*^−^*^4^, 5 *×* 10*^−^*^3^, and 2 nucleotide subs/site, which yielded approximately 80, 800, and 95,000 unique site patterns respectively. The full process is detailed in the methods section below, and the distributions of the number of unique site patterns in each sequence alignment dataset are represented in Fig. 3.

### 4.1 Simulations

#### 4.1.1 Data generation

We scaled the summary trees generated from the empirical analysis to three different total tree lengths (i.e. sum of branch lengths): 4 *×* 10*^−^*^4^, 5 *×* 10*^−^*^3^, and 2 subs/site. This allowed us to simulate the effect of increasing the average number of unique site patterns, without having to increase the sequence length, which would have required considerably more computing storage.

Our first set of simulations assumed a strict molecular clock. To set the tree length, we rescaled the branch lengths using a custom Python script that iterates over the branches and scales the branch rates so that they would add up to the desired new length. Our second set of simulations assumed a relaxed molecular clock with an underlying lognormal distribution (Bromham et al., 2018, Drummond et al., 2006). For this model we employed another Python script to iterate through the branches of each tree, but in this case, we also multiplied them by random draws from a lognormal distribution with *mean* = 2 and *σ* = 1 and then rescaled them to desired total lengths. This final rescaling multiplies every branch rate by the same constant, namely the desired tree length divided by the length of the tree before rescaling. The branch rates therefore remain lognormally distributed, with the same standard deviation on the log scale and hence the same coefficient of variation, and only their mean is shifted. Rescaling thus sets the total tree length without altering the degree of among-lineage rate variation, which is the quantity the relaxed clock is meant to capture. We then used AliSim (Ly-Trong et al., 2023) in IQ-TREE 2 (Minh et al., 2020) to simulate 100 sequence alignments with a length of 200,000 nucleotides for each combination of month and tree length, for the strict molecular clock model case. For the relaxed molecular clock model, we only simulated 20 alignments of each combination. The difference in the number of simulations is because we uncovered consistent results for all simulations under a strict clock model and simply aimed to incorporate a relaxed clock as well, to observe the effects, while using computational resources responsibly and efficiently.

#### 4.1.2 Analysis of simulated data

We analysed all 900 sequence alignments from the strict molecular clock model, and all 180 alignments from the relaxed clock model in BEAST2 v2.7.7 (Bouckaert et al., 2019). As with the inference of the empirical data, we selected a HKY+Γ_4_ substitution model with empirical base frequencies, the matching molecular clock model (strict or relaxed with an underlying lognormal distribution), and an exponential growth coalescent tree prior. We used Markov chain Monte Carlo (MCMC) to sample from the posterior distribution, with a chain length that ensured that the effective sample size (ESS) of each parameter was at least 200 by employing the built-in loganalyser package in BEAST2. We sampled parameters and trees every 10^3^ steps. The priors used for the Bayesian inference are shown in Table 1. The priors for our simulations are intended to be vague while containing the value used to generate the data. For the empirical data we also selected vague priors but their parameterisation represents our expectation of their resulting estimates. For example, the priors on the clock rate for H1N1 influenza virus and HBV are centred on their expected clock rates from previous studies, of the order of 10*^−^*^3^ and 10*^−^*^6^ subs/site/year, respectively (Hedge et al., 2013, Kocher et al., 2021). We fixed the tree topology in all of these analyses, and we computed uncertainty in node times using TreeAnnotator, part of the BEAST2 package.

**Table 1:** Prior configuration for molecular clock model and tree prior. Note that there are two possible molecular clock models, the strict and relaxed molecular clock model with an underlying lognormal distribution, which were used separately to match the simulation conditions. For the standard deviation of the lognormal distribution (relaxed clock), the default prior provided by BEAST2 was used.

| Parameter | Prior |
| --- | --- |
| Scaled population size, $\Phi$ | *Exponential*(*mean* = 1,000) |
| Exponential growth rate, $r$ | *Laplace*(*mean* = 0, *scale* = 50) |
| Evol. rate (strict clock) | $\Gamma$(*shape* = 0.5, *rate* = 1) |
| Mean of the lognormal (relaxed clock) | $\Gamma$(*shape* = 0.5, *rate* = 1) |
| Standard deviation of the lognormal (relaxed clock) | $\Gamma$(*shape* = 0.54, *rate* = 0.38) |

Each of our simulation replicates resulted in a posterior distribution of parameters and trees. To explore the infinite-sites behaviour we obtained the HPD intervals of nodes by summarising the heights of the trees sampled from the posterior. Note that in our simulations, the tree topology is fixed to the truth, such that all trees in the posterior contain the same set of nodes and thus their heights are a continuous distribution for which we can compute mean values and HPDs. This is a necessary step to study uncertainty in divergence time estimates independently of phylogenetic uncertainty.

We fit linear regressions using ordinary least squares, as implemented in the Python library Scipy (Virtanen et al., 2020), of the HPD interval width of internal nodes as a function of their mean distance to the closest tip-calibration. In this context, the slope of the regression is the rate at which uncertainty increases, the y-intercept is the expected uncertainty of a node with distance of zero to its closest tip-calibration, and the extent to which the points deviate from the line of best fit is a measure of their infinite-sites behaviour. To assess the latter we chose to use the root mean squared error (RMSE) as a metric of linearity instead of *R*^2^, because the RMSE has the same units as the dependent variable (e.g. years), whereas *R*^2^ has no units. In our comparison the units are relevant because they provide a straightforward understanding of uncertainty. Moreover, the RMSE quantifies the typical magnitude of the deviation between prediction and truth in the units of the dependent variable, whereas *R*^2^ is normalised, which is problematic for very small values, as we see here.

#### 4.1.3 Empirical data

We used samples taken from the 2009 North American H1N1 influenza virus, collected by Hedge et al. (2013), from the months June, August, and December to provide insight into the effect of increasing the number of taxa on our study of the infinite-sites theory (the number of taxa and unique site patterns for each month are given in Fig. S1 of the Supplemental Material). The datasets provided in the original paper for these three months represent data taken from January to June, January to August, and January to December, which we refer to as June, August, and December respectively in this manuscript. We also analysed a dataset of 100 HBV genomes collected by Kocher et al. (2021) and curated by Weber et al. (2025). We set the same Bayesian phylogenetic model as for our simulations and the priors described in Table 2. We analysed our empirical data using BEAST2, by running the MCMC chains until all model parameters had reached an ESS of at least 200 as in our simulations.

**Table 2:**
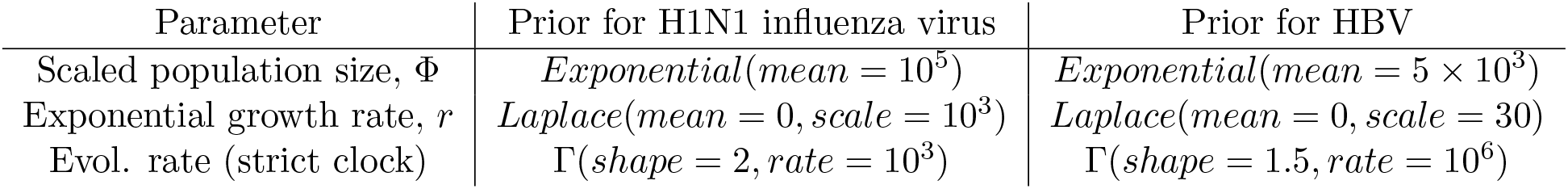
Prior configuration for molecular clock model and tree prior. The substitution model configuration was described above.

| Parameter | Prior for H1N1 influenza virus | Prior for HBV |
| --- | --- | --- |
| Scaled population size, $\Phi$ | *Exponential*(*mean* = $10^5$) | *Exponential*(*mean* = $5 \times 10^3$) |
| Exponential growth rate, $r$ | *Laplace*(*mean* = 0, *scale* = $10^3$) | *Laplace*(*mean* = 0, *scale* = 30) |
| Evol. rate (strict clock) | $\Gamma$(*shape* = 2, *rate* = $10^3$) | $\Gamma$(*shape* = 1.5, *rate* = $10^6$) |

## 5 Data availability

All empirical data used in this study are available on GenBank and were published in a previous study by Hedge et al. (2013). Computer code, analysis files, and sequence datasets in this study are available at: https://github.com/simonaivanov/pasteur_infinite_sites_paper_clean

## 6 Competing interests

None.

## 7 Acknowledgements

We are grateful to Prof. Laura Katz, Prof. Beatriz Mello, Prof. Jeffrey Thorne, and two anonymous reviewers for their useful feedback on earlier versions of this manuscript.

## 8 Supplemental Material

See supplementary material online.

## 9 Funding

This work received funding from the Inception program (Investissement d’Avenir grant ANR-16-CONV-0005 awarded to SD) and from a project grant from the Agence Nationale de Recherche AAPG2024 (project TrAM awarded to SD).

## Notes

### Competing Interest Statement

The authors have declared no competing interest.

### Summary of Updates

Additional simulation study to understand the calibration information from closest tips, relative to closest sister. Described in detail in supp info and in the main text.

