## Supplemental Material for "The limits of Bayesian estimates of divergence times in measurably evolving populations"

| Month | Taxa | Unique site patterns |
|---|---|---|
| April | 34 | 131 |
| May | 100 | 345 |
| June | 164 | 582 |
| July | 186 | 692 |
| August | 206 | 807 |
| September | 243 | 1006 |
| October | 276 | 1186 |
| November | 307 | 1370 |
| December | 328 | 1526 |

Figure S1: Table describing the number of taxa and unique site patterns for each month of the original 2009 H1N1 Influenza virus dataset provided by Hedge et al.

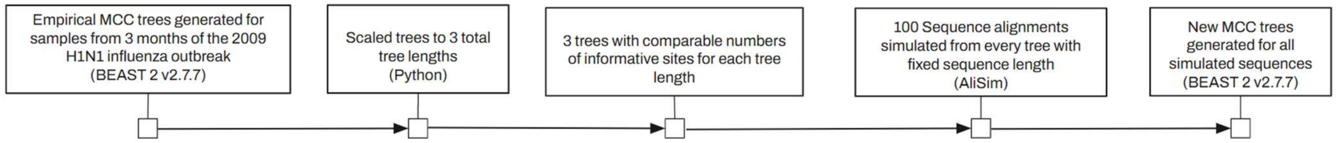


Figure S2: Schematic explaining the steps used to create the simulated trees.

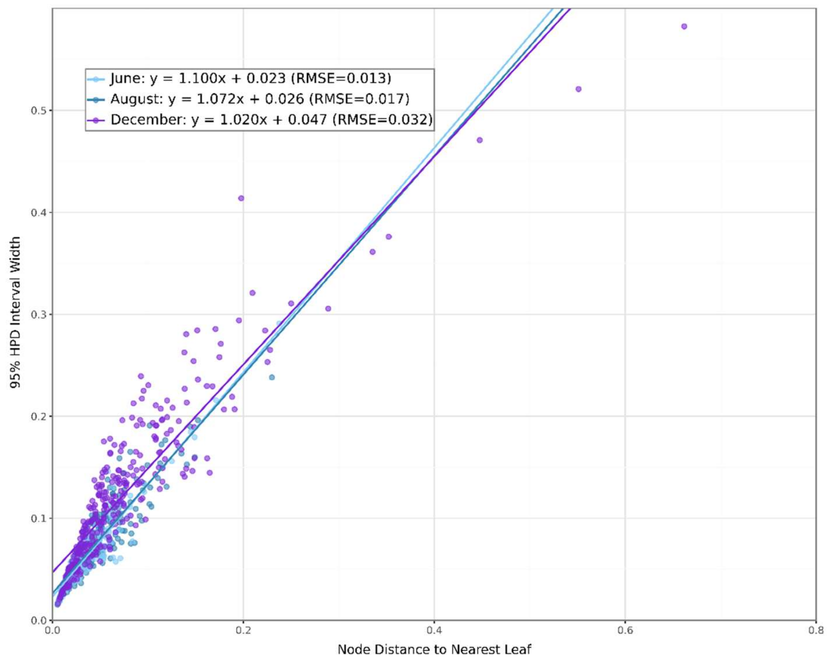


Figure S3: Infinite sites plot for tree length of 0.0004 subs / site, when the distance to closest leaf node is taken to be the minimum of the distance to the node of interest's closest leaf node and that of its parent.

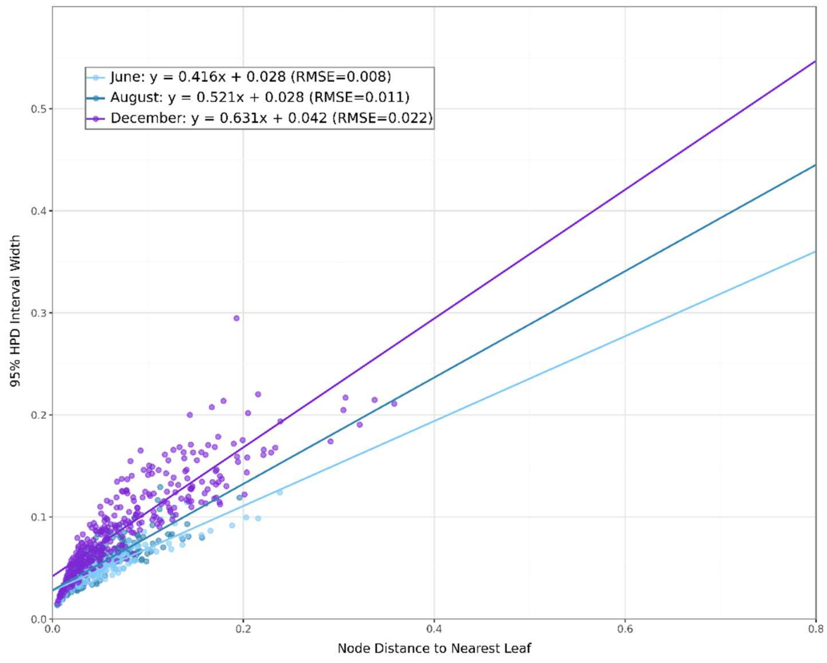


Figure S4: Infinite sites plot for tree length of 0.005 subs / site, when the distance to closest leaf node is taken to be the minimum of the distance to the node of interest's closest leaf node and that of its parent.

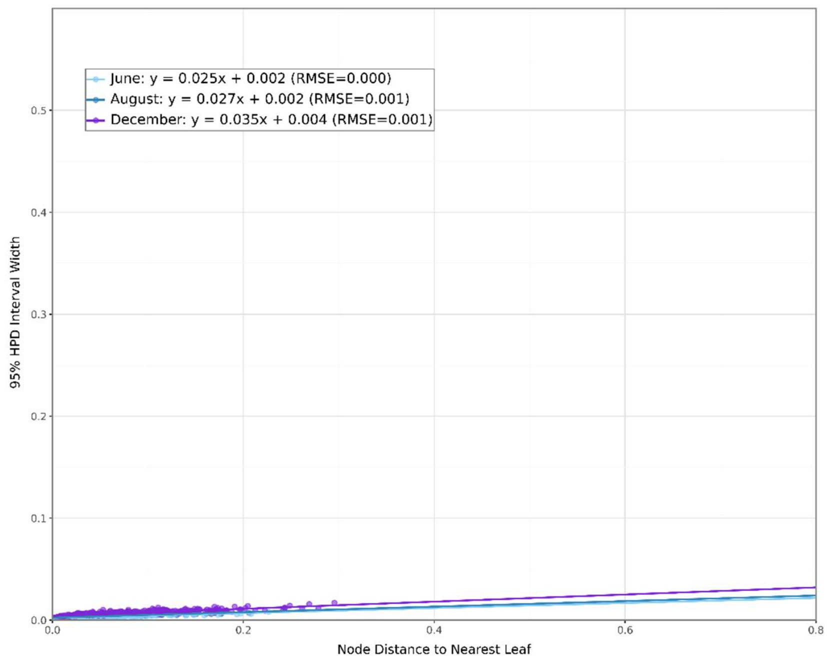


Figure S5: Infinite sites plot for tree length of 2 subs / site, when the distance to closest leaf node is taken to be the minimum of the distance to the node of interest's closest leaf node and that of its parent.

### 1 Calibration information in heterochronous data

In this section we assess the degree to which the closest descendant tips and the closest sister tips provide calibration information for internal nodes. The key question is whether nodes with very long distances to their closest descendant tip, but for which the closest sister tip distance is short, display lower uncertainty. In other words, *when the sister distance is shorter than the distance to the descendant tip, is the uncertainty lower than in the opposite case?*

For a subset of our data we conduct infinite sites plots using the closest descendant tip (as in the rest of our study) or by taking the closest sister tip when its distance to the node in question is shorter than that to the closest descendant.

In fig. S6 we illustrate a hypothetical scenario where a sister tip sits closer to a node of interest and may provide more calibration information than a descendant tip. Here we are interested in node $e$, for which node $a$ is the 'tip-calibration' because it is the descendant node with the shortest distance (in units of time) to node $e$, such that it should provide most of the calibration information. However, in this example, the temporal distance from $e$ to $a$ is 20 units of time, whereas its closest sister tip, $c$ has a distance of only 5 time units and is thus much closer in time and is potentially more informative than $a$. In molecular clock analyses of heterochronous data, all sampling times provide calibration information (to the extent that they are provided with low uncertainty), but here, nodes $a$ and $c$ are expected to provide different amounts of information for the estimate of the age of $e$.

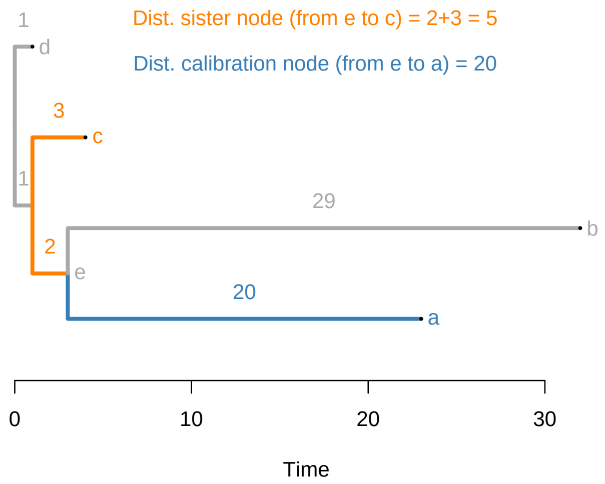


Figure S6: Example of a heterochronous phylogenetic tree and sources of calibration.

For our analyses of data sampled until the month of April and using four tree lengths, we find that the distance to sister tips is shorter than the standard tip-calibration only a small number of times (fig. S7; for two nodes when the tree length was 0.003 or 0.012, and up to three nodes when the tree length was set to 0.006 or 0.024). The regressions that included these points have higher slopes, implying a *departure* from infinite sites behaviour in the sense that our most informative simulations had slopes that were flat and low RMSE. This is to be expected, however, because their distance is by definition shorter on the $x-$axis.

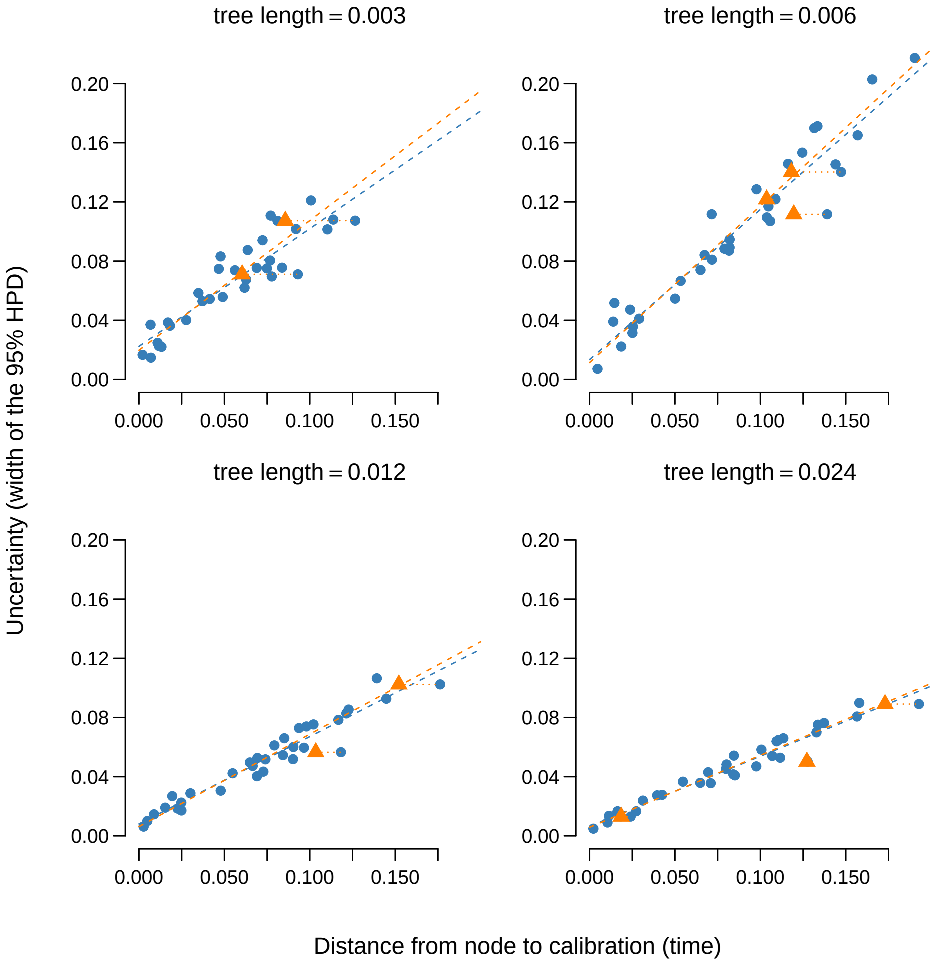


Figure S7: Uncertainty in internal node ages, defined as the width of the 95% HPD interval, as a function of the distance to the calibration for four tree lengths of data sets simulated under the influenza virus data set sampled until the month of April. Blue points are where we consider the closest descendant tip as the calibration (i.e. the 'tip-calibration' in the main text). For internal nodes for which the closest sister tip is closer in distance to the node in question than the tip-calibration, we also show the corresponding distance to the closest sister tip in the orange triangles. The dotted lines connect the closest sister tip to the tip-calibration (the descendant) at constant $y$, so the two points joined by a line belong to the same node. The dashed lines show the least-squares regression.

In the case of the data simulated under the influenza virus tree sampled up to the month of December, where there were many more samples in total, we observed many more instances where the closest descendant tip has a longer distance to the node in question than the closest sister (fig S8). Here, the regression slopes are also larger than those when using only the tip-calibrations.

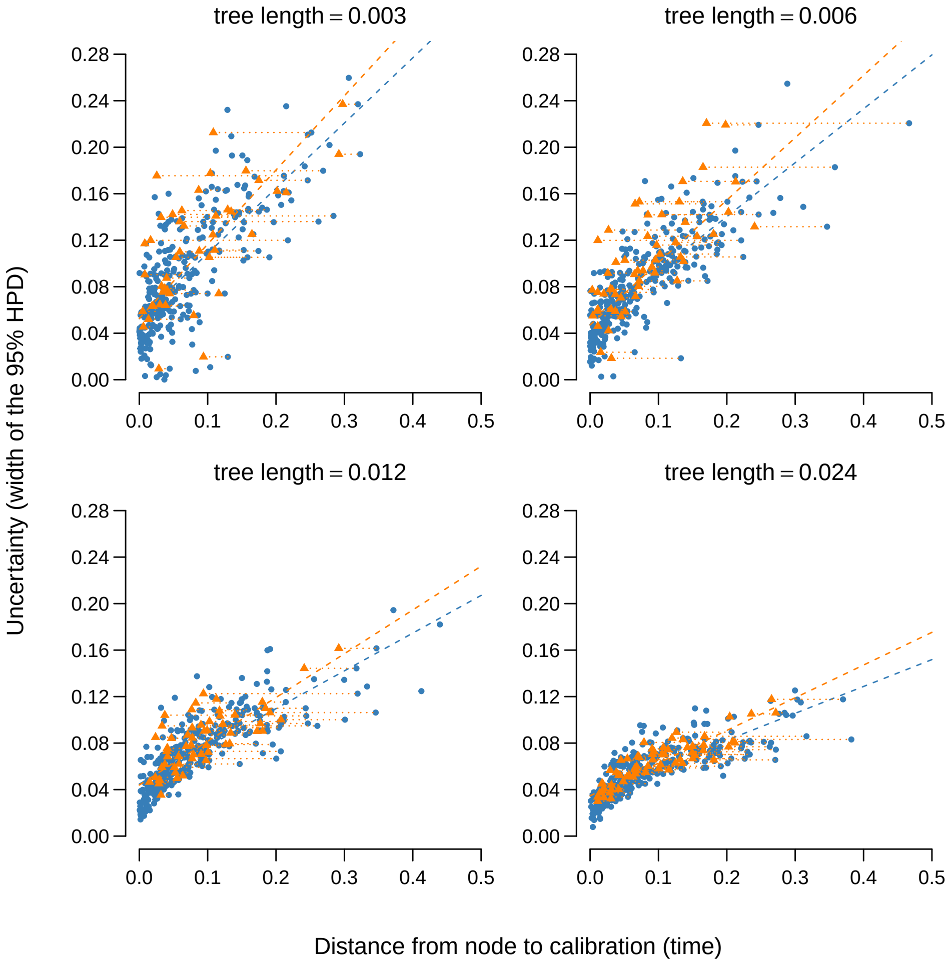


Figure S8: Uncertainty in internal node ages, defined as the width of the 95% HPD interval, as a function of the distance to the calibration for four tree lengths of data sets simulated under the influenza virus data set sampled until the month of December. The colours and symbols match those of fig S7.

We also show scatter-plots of the uncertainty in node age versus the difference in distance between the tip-calibration (closest descendant tip) and the closest sister tip. We do not find an obvious association between internal node uncertainty and the magnitude of this difference (fig S9); the comparison between the two classes of node themselves is made below.

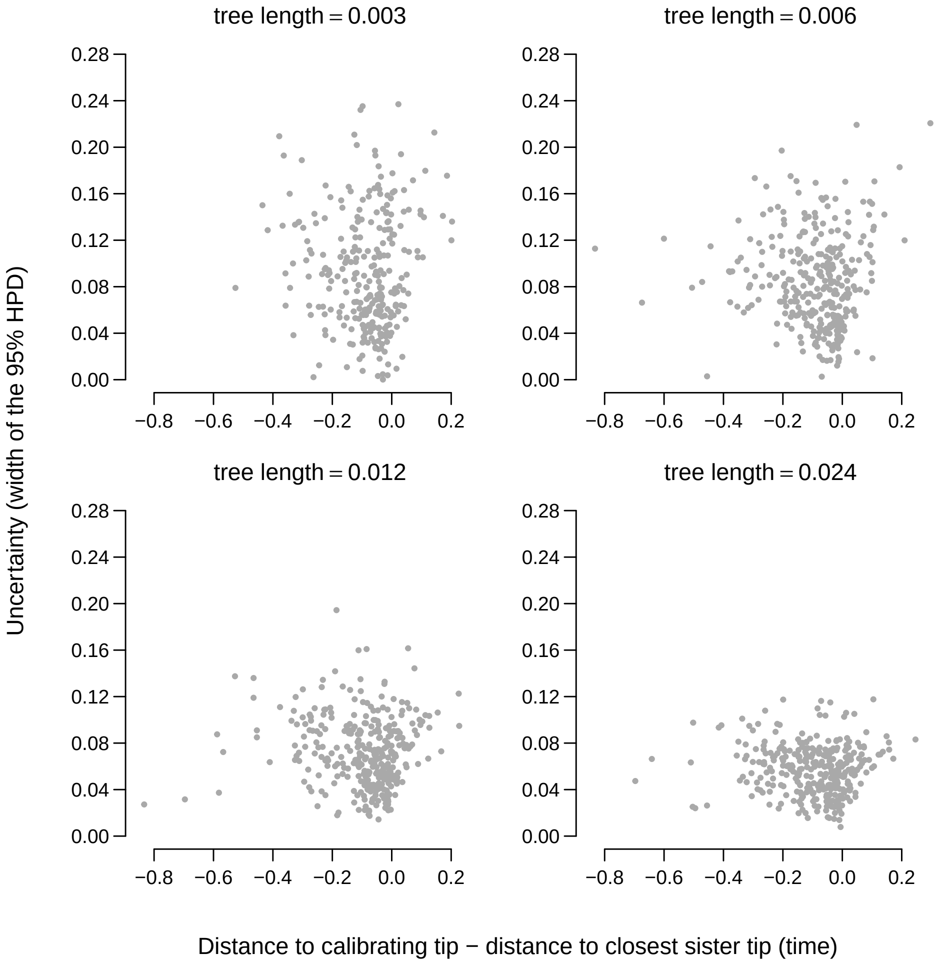


Figure S9: Uncertainty in internal node ages versus the difference between two potential calibrations (the tip–calibration or the sister tip). Negative values on the x axis mean the descendant tip of a node is the nearer of the two.

To compare the overall uncertainty of nodes for which the tip-calibration was closer than the closest sister tip, we plotted the uncertainty for two sets of nodes (figs S10 and S11). Although there is a broad overlap in the distributions of points along the $y$ axis, we note that the mean uncertainty tends to be higher in nodes for which the closest sister is closer than the closest descendant. This comparison is not made at matched distances, however. A node falls into that class only when its own descendants are distant in time, so the class is enriched for long distances to the closest descendant, and a higher mean uncertainty is expected on those grounds alone. The experiments in the next section compare the two kinds of calibration at matched distances, which is what this question requires.

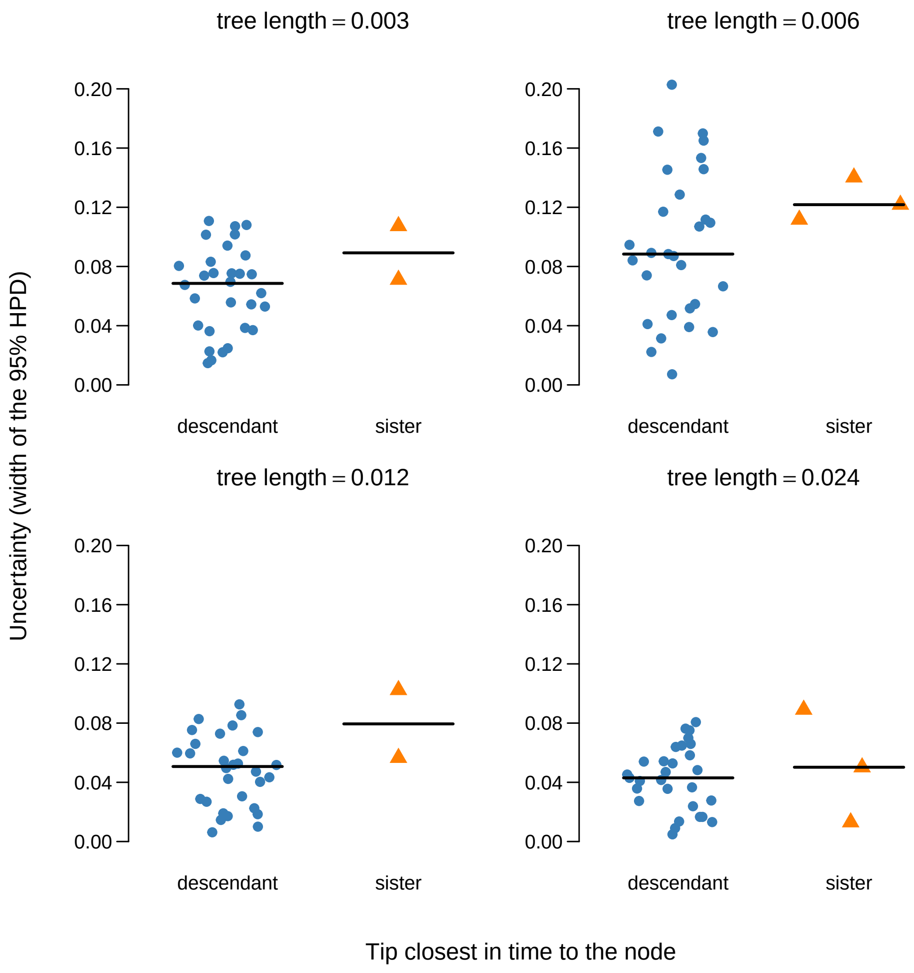


Figure S10: Uncertainty in internal node ages for two classes of node, under four tree lengths, for data simulated under the influenza virus tree sampled until the month of April. Nodes are split according to whether the tip closest in time is one of their own descendants (blue circles) or sits on the sister side of the parent (orange triangles). The horizontal spread within each group is jitter proportional to the local density of the points and carries no further meaning; the heavy horizontal bar is the group median.

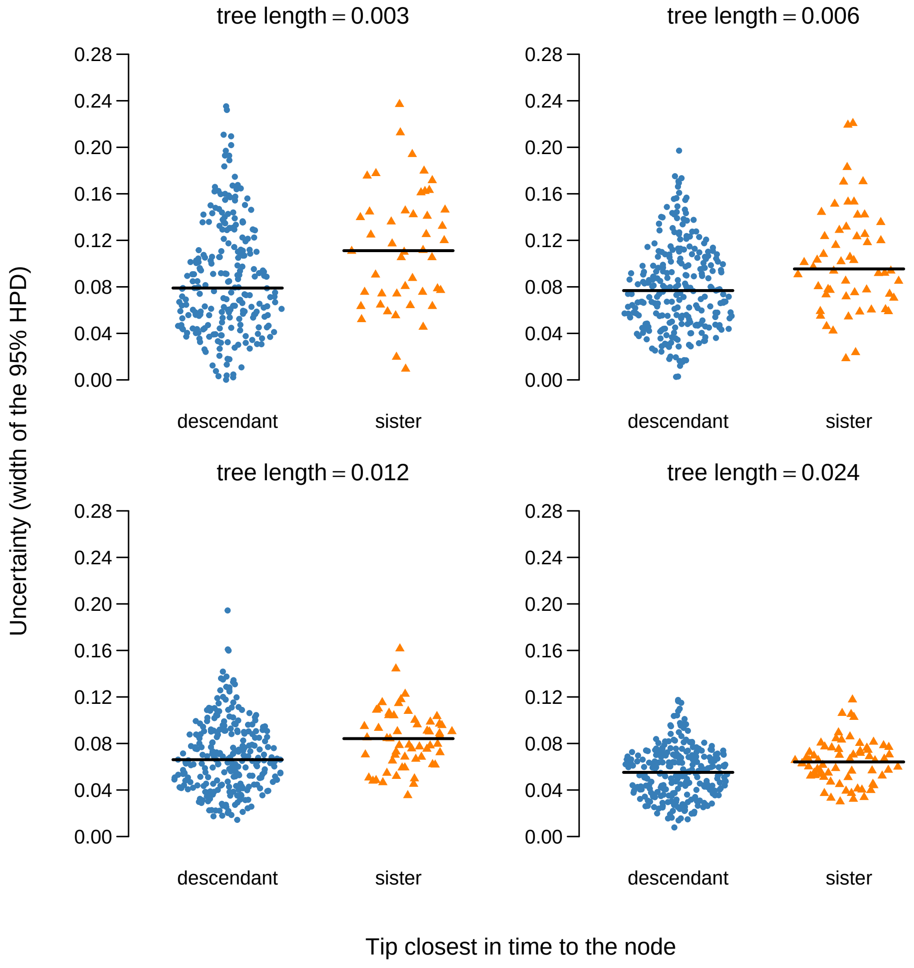


Figure S11: As in fig S10, but for data simulated under the influenza virus tree sampled until the month of December, which contains many more nodes whose closest tip in time is not a descendant.

#### 2 Disentangling sources of calibration information

We conducted an additional set of simulation experiments to investigate the information from both kinds of calibration information. We took the phylogenetic tree topology from fig S6, which only contains four tips. To define the branch lengths in units of time we sampled numbers from an exponential distribution with mean 1 and assigned them to the six branches, ensuring that the largest values were those for the terminal branches leading up to taxa $a$ and $b$. As a result, the distance from node $e$ to its closest sister tip, $c$ tended to be shorter than that to the closest descendant ($a$ or $b$). To exacerbate this pattern, we multiplied the length of the two terminal branches leading up to $a$ and $b$ by two. We simulated sequence alignments of length 1,000 nucleotides under a JC substitution model and assuming a strict molecular clock with rate 0.01 subs/site/time. We generated 100 of such alignments, which had between 28 and 174 variable sites.

We analysed each data set under a fixed tree topology, with the sampling times as calibrations, and we matched the generating process as closely as possible. That is, a JC substitution model and a strict molecular clock. For the tree prior we assumed a constant-size coalescent, which is not the matching generating process, but we chose it for statistical simplicity. For the population size we chose an exponential distribution with mean 10 as a prior, while for the molecular clock rate we used a Gamma distribution with alpha=2 and beta=200 (for a mean of 0.01, the true value).

We sampled from the posterior distribution with BEAST2 (Bouckaert et al., 2019) as in our Methods (in the main text). Because these simulations consisted of very small data sets, we set the chain length to $10^7$, sampling every 5,000 steps, which resulted in effective sample sizes of over 1,000 for all parameters.

We found that the slope of the uncertainty of node $e$ as a function of the distance to the calibration was steeper when we considered the sister tip than the closest descendant, and that the RMSE for the sister tip was the lower of the two (fig S12).

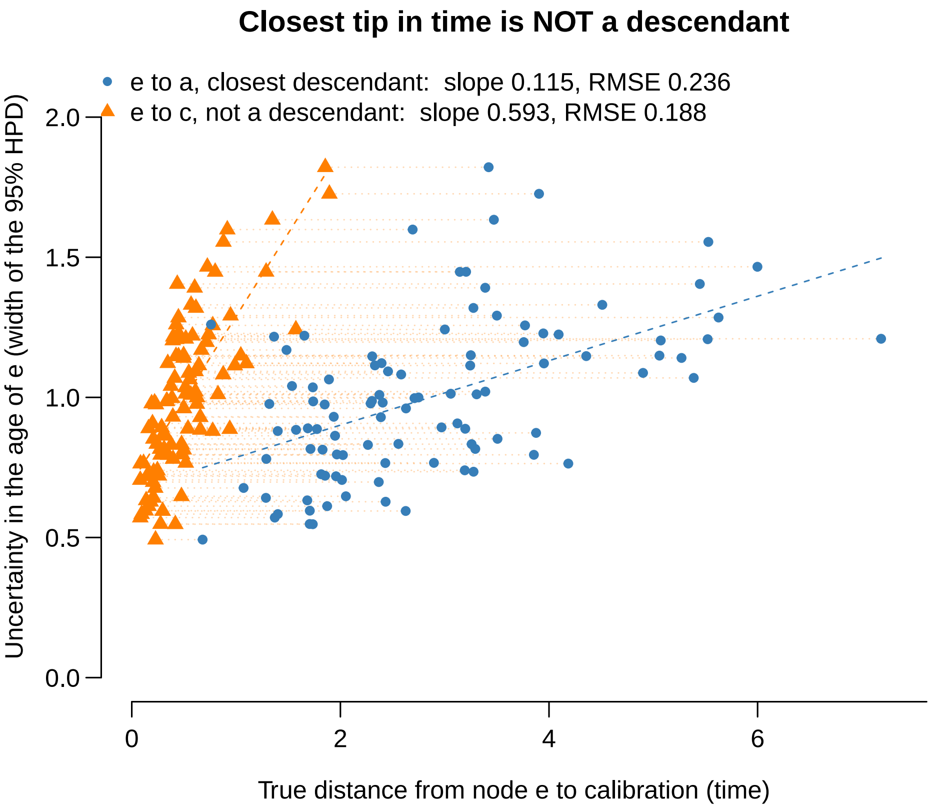


Figure S12: Uncertainty as a function of distance to calibrating tips for simulations under conditions that resemble those of tree in fig S6. The colours and shapes match those of fig S10. However, here the points correspond to the node $e$ for 100 independent simulation replicates. The dotted lines are used to connect the same data set for the two calibration measurements ($e$ to $a$ or $e$ to $c$).

It is tempting to conclude that uncertainty decreases faster relative to the closest sister tip, but it is important to note that, by design, these distances are necessarily shorter than those of the closest descendant tip. In fact, it is equivalent to compressing the points in blue along the $x$-axis in fig S12.

We decided to perform an additional set of simulations with the same tree topology as in fig S6, but we sorted the branch lengths so that the descendant tips from $e$ had shorter distances than the sister tip $c$ (fig S13). These two simulation experiments set the scene for a more fair comparison. In the first set we effectively vary the length of the path between $e$ and $c$ when it is always shorter than that from $e$ to $a$. In the second set, we vary the distance from $e$ to $b$, while ensuring that it is shorter than from $e$ to $c$.

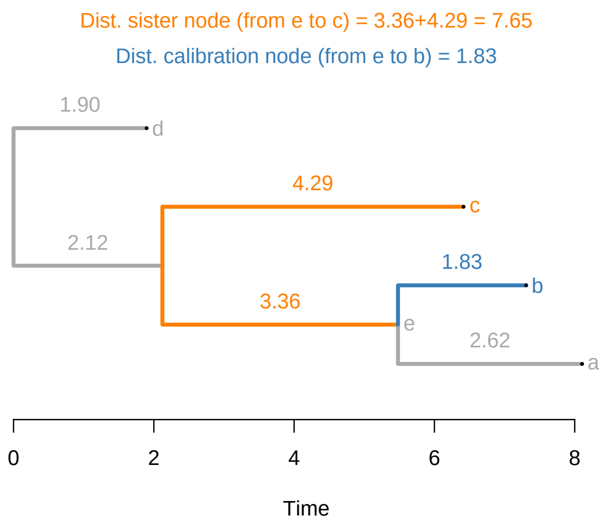


Figure S13: Example of a heterochronous phylogenetic tree and sources of calibration. In contrast to fig S6 tip $b$ is much closer to node $e$ and is thus expected to provide most calibration information.

This experiment revealed two important points. First, when the closest tip is a descendant the uncertainty can attain much lower values, sometimes approaching zero (fig S14). Second, if we consider the slope of the infinite sites plot as an indication of the amount of information conveyed by the calibration, then descendant tips are potentially more informative than sister tips here, given that they yield a much steeper slope over the same range of distances (1.41 against 0.08).

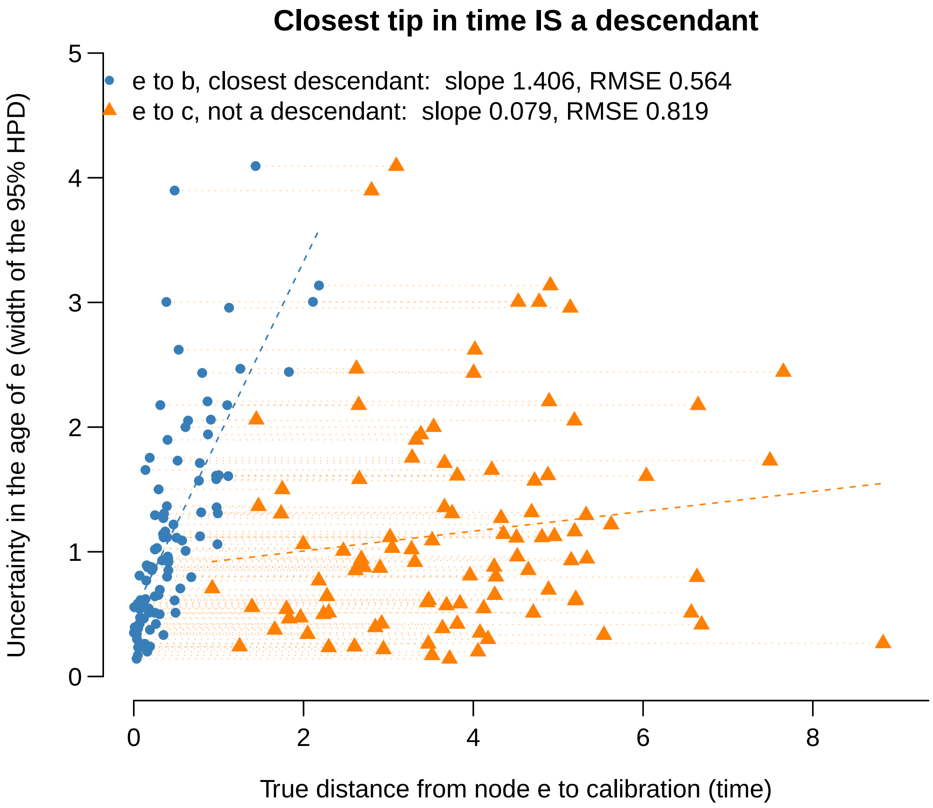


Figure S14: Uncertainty as a function of distance to calibrating tips for simulations under conditions that resemble those of tree in fig S13. The colours and shapes match those of fig S12.

For a more controlled set of simulation experiments we considered a fixed tree where we only varied the length of the path between $e$ and $c$, as shown in fig S15. We considered ten sets of values, ranging from a total of 0.1 units of time to 1, which is the same as the distance to the closest tip $b$. Then we devised a similar procedure in which we only vary the distance to the closest descendant (see fig S16). Concretely, we generated 10 trees in each case, with varying distance to the calibration of interest, with 10 replicates of each.

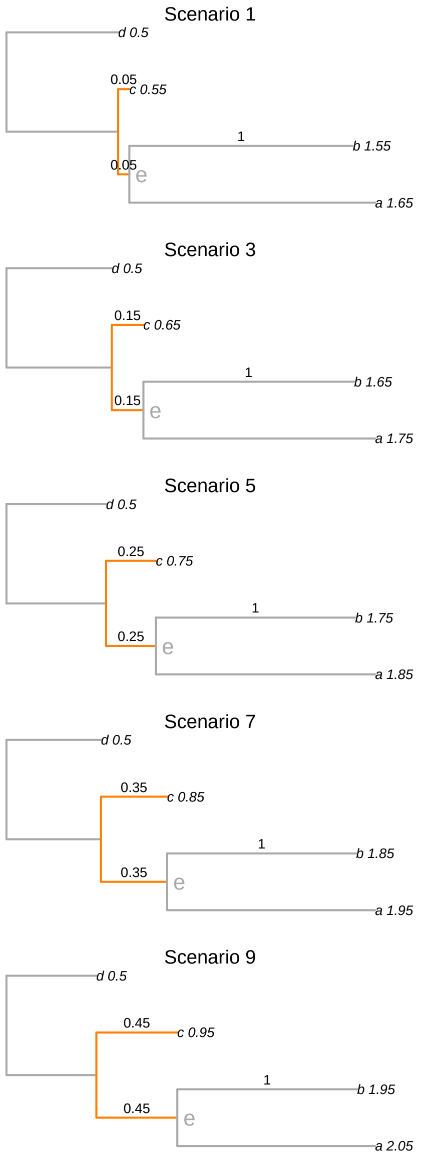


Figure S15: Phylogenetic time trees where we vary the distance to the sister tip only (relative to node $e$). Here the tips show their corresponding sampling times.

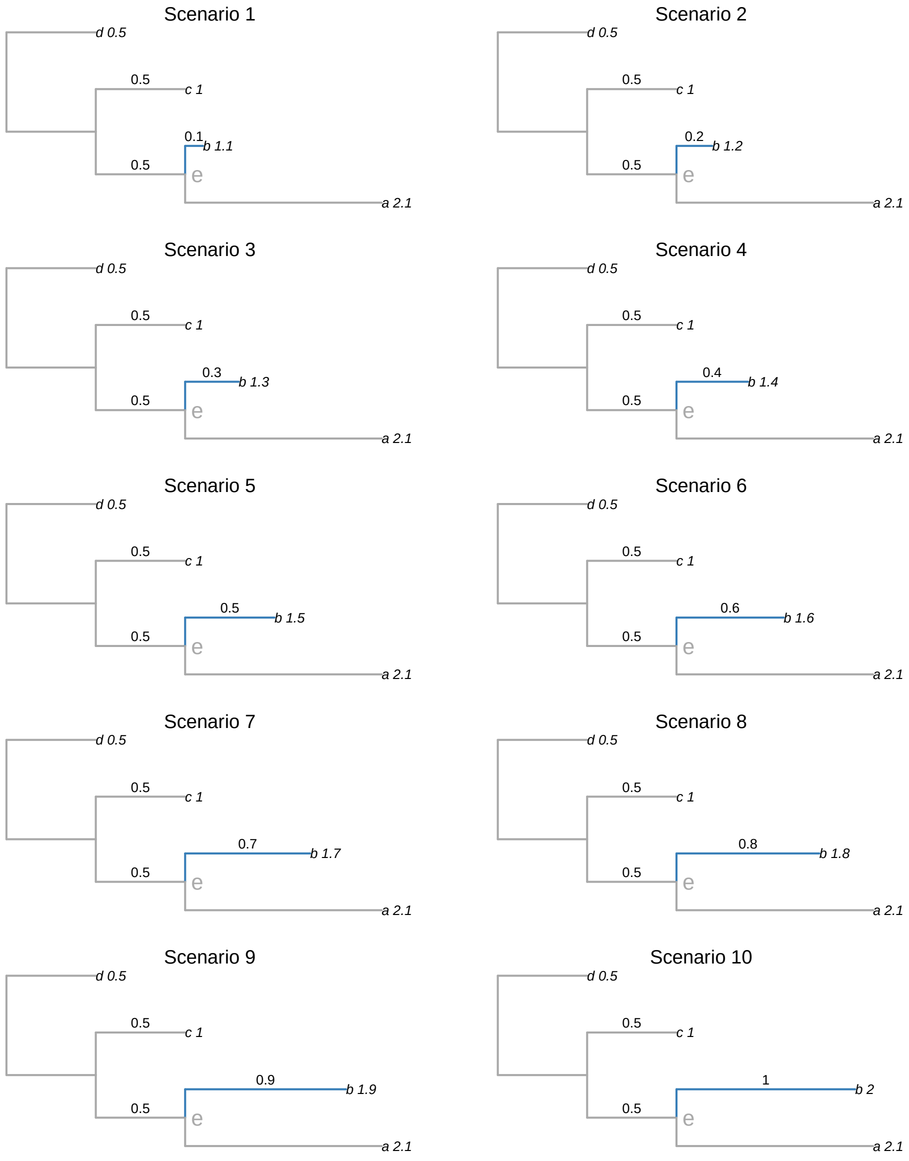


Figure S16: Phylogenetic time trees where we vary the distance to the closest descendant tip only (relative to node $e$). Here the tips show their corresponding sampling times.

A regression of the uncertainty as a function of the true distance from node $e$ to both calibration points revealed very similar behaviour (fig S17). The slope for the distance to the descendant tip is somewhat steeper than that for the sister tip, while the RMSE is smaller for the latter. Although this finding implies a stronger effect size for the descendant tip, meaning that it is more informative, we note that the differences are small (e.g. 0.728 for the descendant, compared to 0.68 for the sister tip).

The sum of these simulation experiments demonstrates that both the closest descendant *and* the closest sister tips provide calibration information to a given internal node. In our main study, we chose to use the descendant tip as the 'calibration-tip'. Indeed, it is the most common pattern found in empirical data (see figs S7 and S8), and that which is most easily generated by coalescent and birth-death processes (see Möller et al., 2018). When there exists a sister tip that is much closer than the closest descendant, it may convey more calibration information. In particular, compare the pattern of fig S12 to that of fig S14, where the order of the slopes and RMSE are the opposite. For practical applications it is advisable to compare infinite site plots that consider the closest descendant tip and the closest sister tip to explain uncertainty in node times.

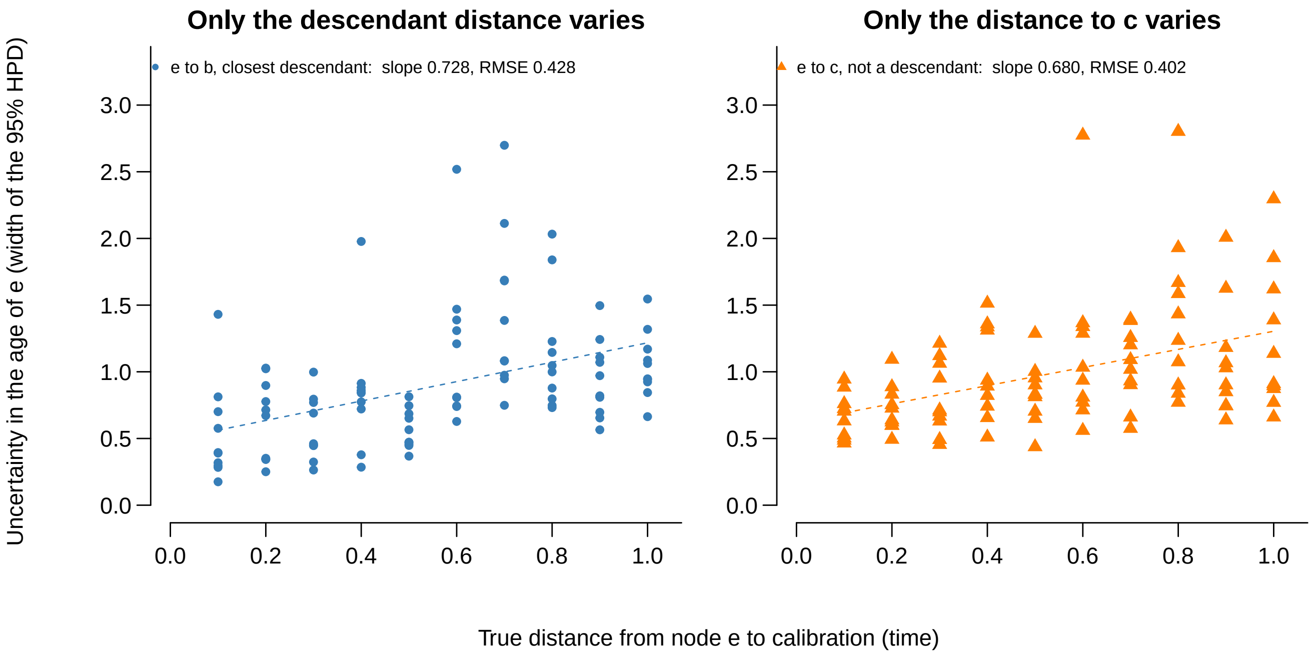


Figure S17: Regressions of the uncertainty in node $e$ as a function of its distance to the closest calibration. In the left panel we vary the distance to the descendant tip (fig S16), while the panel on the right denotes the situation where the distance to the sister tip varies (fig S15).
